# Magnetic nanocomplexes coupled with an external magnetic field modulate macrophage phenotype – a non-invasive strategy for bone regeneration

**DOI:** 10.1101/2023.09.02.556050

**Authors:** Harshini Suresh Kumar, Zhongchao Yi, Sheng Tong, Ramkumar T. Annamalai

## Abstract

Chronic inflammation is a major cause for the pathogenesis of musculoskeletal diseases such as fragility fracture, and nonunion. Studies have shown that modulating the immune phenotype of macrophages from proinflammatory to prohealing mode can heal recalcitrant bone defects. Current therapeutic strategies predominantly apply biochemical cues, which often lack target specificity and controlling their release kinetics *in vivo* is challenging spatially and temporally. We show a magnetic iron-oxide nanocomplexes (MNC)-based strategy to resolve chronic inflammation in the context of promoting fracture healing. MNC internalized pro-inflammatory macrophages, when coupled with an external magnetic field, exert an intracellular magnetic force on the cytoskeleton, which promotes a prohealing phenotype switch. Mechanistically, the intracellular magnetic force perturbs actin polymerization, thereby significantly reducing nuclear to cytoplasm redistribution of MRTF-A and HDAC3, major drivers of inflammatory and osteogenic gene expressions. This significantly reduces *Nos2* gene expression and subsequently downregulates the inflammatory response, as confirmed by quantitative PCR analysis. These findings are a proof of concept to develop MNC-based resolution-centric therapeutic intervention to direct macrophage phenotype and function towards healing and can be translated either to supplement or replace the currently used anti-inflammatory therapies for fracture healing.

## 1. Introduction

Persistent inflammation is a crucial cause for the pathogenesis of musculoskeletal diseases such as fragility fracture, nonunion, arthritis, and other rheumatic conditions. Over 10 million individuals suffer from recalcitrant fractures each year in the US. Arthritis (doctor-diagnosed), on the other hand, affected an average of 54.4 million adults between 2013 and 2015, or 23 in 100 adults^1^. Additionally, chronic wounds due to fragility fractures affecting 25% of men and 50% of women over age 50 are a significant cause for pain and disability. The healthcare cost for these conditions is estimated to be over 25 billion dollars by 2025^2^. Currently, there are no reliable first-line therapies that stimulate healthy bone tissue formation and promote functional recovery.

Bone homeostasis is biologically entangled with the immune response, where macrophage plays an indispensable role in maintaining bone integrity and metabolism. Macrophages are a heterogeneous group of immune cells that adopt discrete functional phenotypes depending on their microenvironment^3^. Osteomacs, a type of tissue-resident macrophages, play a significant role in osteoblast mineralization *in vitro* and induce osteoblast differentiation *in vivo*^4, 5^. Further, studies in the tibial fracture model show that osteomacs promote collagen I deposition and mineralization in response to bone damage^5, 6^. Cytokines secreted by macrophages, such as interleukin-1 (IL-1) and interleukin-6 (IL-6), are also shown to promote angiogenesis and primary cartilaginous callus formation, respectively^7^. Although acute inflammation is crucial for bone regeneration and homeostasis^8, 9^, a prolonged inflammation may lead to suppressed healing^10-13^ and bone resorption^14^.

Macrophages are also implicated in the pathogenesis of inflammatory musculoskeletal conditions, including rheumatoid arthritis (RA)^15, 16^, osteoarthritis (OA)^17^, ankylosing spondylitis (AS)^18^, and peri-implant osteolysis (PIO)^19^. In arthritis, macrophages are the primary source of proinflammatory cytokines causing synovial inflammation and subsequent osteoclastogenesis leading to joint erosion and damage^15-17^. Clinical data also show a strong correlation between the accumulation of CD68^+^ macrophages and the extent of joint destruction in RA patients^20^. Likewise, the quantity and activation state of the macrophages for AS patients are correlated to disease severity^21^. In addition, the cytokines secreted by macrophages can induce Th1 and Th17 differentiation, further propagating the tissue damage^22^. In PIO and wear particles, macrophages are present abundantly in the granulomatous periprosthetic membrane. The wear particles trigger proliferation, differentiation, and activation of macrophages^19^ and stimulate them to produce chemokines that promote osteoclastogenesis and bone resorption^23^.

Studies have shown that modulating the immune phenotype of macrophages from proinflammatory to prohealing mode can heal recalcitrant bone defects and rheumatic ailments^24-26^. They mainly apply biochemical factors that predominately work to suppress proinflammatory cytokines or incite an M2-like phenotype. The other facets of tissue healing, such as ECM deposition, angiogenesis, and osteogenesis, are largely overlooked. In addition, biochemical interventions lack target specificity, and controlling their release kinetics *in vivo* is challenging spatially and temporally. Alternatively, immunomodulation through physical interventions such as modifying topography^27^, spatial confinement^28^, stiffness^29^, and porosity^30^ of a provisional matrix could reduce systemic effects. But the poor understanding of macrophage mechanobiology and the long-term progression of these physical factors makes them clinically less attractive. Hence the clinical transition of the immunomodulatory strategy requires cell-targeting capability while allowing precise steering of the inflammatory response. Here we describe a unique approach using magnetic nanocomplexes (MNC) to regulate macrophage immunophenotype by manipulating their cytoskeletal dynamics. These MNC are readily customizable to target desired cells and made from clinically translatable nanomaterials commonly used in tumor imaging and thermal therapy^31^. MNC are readily internalized by macrophages through endocytosis and accrued in lysosomes. Then using an external magnetic field, an intracellular magnetic force can be exerted on the cytoskeleton to manipulate gene expression profiles.

The cytoskeletal dynamics of macrophages are intricately linked to their inflammatory response. Like most adherent cells, macrophages respond to physical cues through actin-cytoskeletal reorganization^32^, nuclear deformation^33^, and subsequent gene expression^34^. During proinflammatory activation via TLR4 signaling, macrophages change their morphology through actin polymerization and subsequent upregulation of several proinflammatory genes, including cyclooxygenase-2, tumor necrosis factor-α (TNF-α), and iNOS^28^. Cyclic straining of macrophage cytoskeleton suppresses proinflammatory cytokines while promoting a pro-reparative or prohealing phenotype^35-37^. Remarkably, mere manipulation of the cytoskeleton using physical cues, without any exogenous factors, is shown to transform macrophage phenotype^28, 38^. This phenotype modulation is mainly attributed to the nuclear translocation of the transcription factor MRTF-A, bound to monomeric G-actin and released upon F actin polymerization^28^. MRTF-Amediated inflammatory gene expression also requires chromatin remodeling and recruitment of transcription initiation machinery. Evidently, cytoskeletal dynamics also govern the spatial redistribution of HDAC3, a histone deacetylase responsible for chromatin remodeling, via IκB-α^39^. Nuclear localization of HDAC3 has been shown to upregulate inflammatory genes, including IL-6, IL-1b, CXCL9, and iNOS in macrophages^28, 40^. Hence, changes in chromatin compaction caused by cytoplasmic-to-nuclear redistribution of HDAC3 during cytoskeletal modulation further tunes macrophage gene-expression programs. Together, actin cytoskeletal dynamics play a significant role in determining macrophages’ transcription machinery and subsequent immunophenotype.

We hypothesized that intracellular magnetic force could elicit transcriptional control of macrophage phenotype and promote fracture healing via MRTF-A release and HDAC3 redistribution **(Fig 1)**. Our approach involves delivering MNC that targets the macrophages at the injury site and, when subjected to an external magnetic field, the intracellular MNC exerts magnetic force to the cytoskeleton, altering cellular phenotype. We show that the MNC can manipulate macrophages phenotype by exerting an intracellular force of 2.27 nN, sufficient to modulate the cytoskeletal dynamics. The size of MNC can be customized and readily conjugated with various surface ligands to facilitate cellular uptake by binding to macrophage cell surface receptors and carrying out specific functions, including cytoskeletal modifications. MNCs were preferentially internalized by pro-inflammatory macrophages and when subjected to external magnetic field, the intracellular magnetic force modified actin cytoskeletal dynamics and restricted nuclear translocation of key transcriptions factors, MRTF-A and HDAC3. Our study shows that MNC-mediated intracellular force modulates macrophage functional phenotype by altering protein and gene expression that are associated to cytoskeleton and promotes pro-healing phenotype. Further, we performed mechanistic studies to validate our findings.

**Figure 1:**
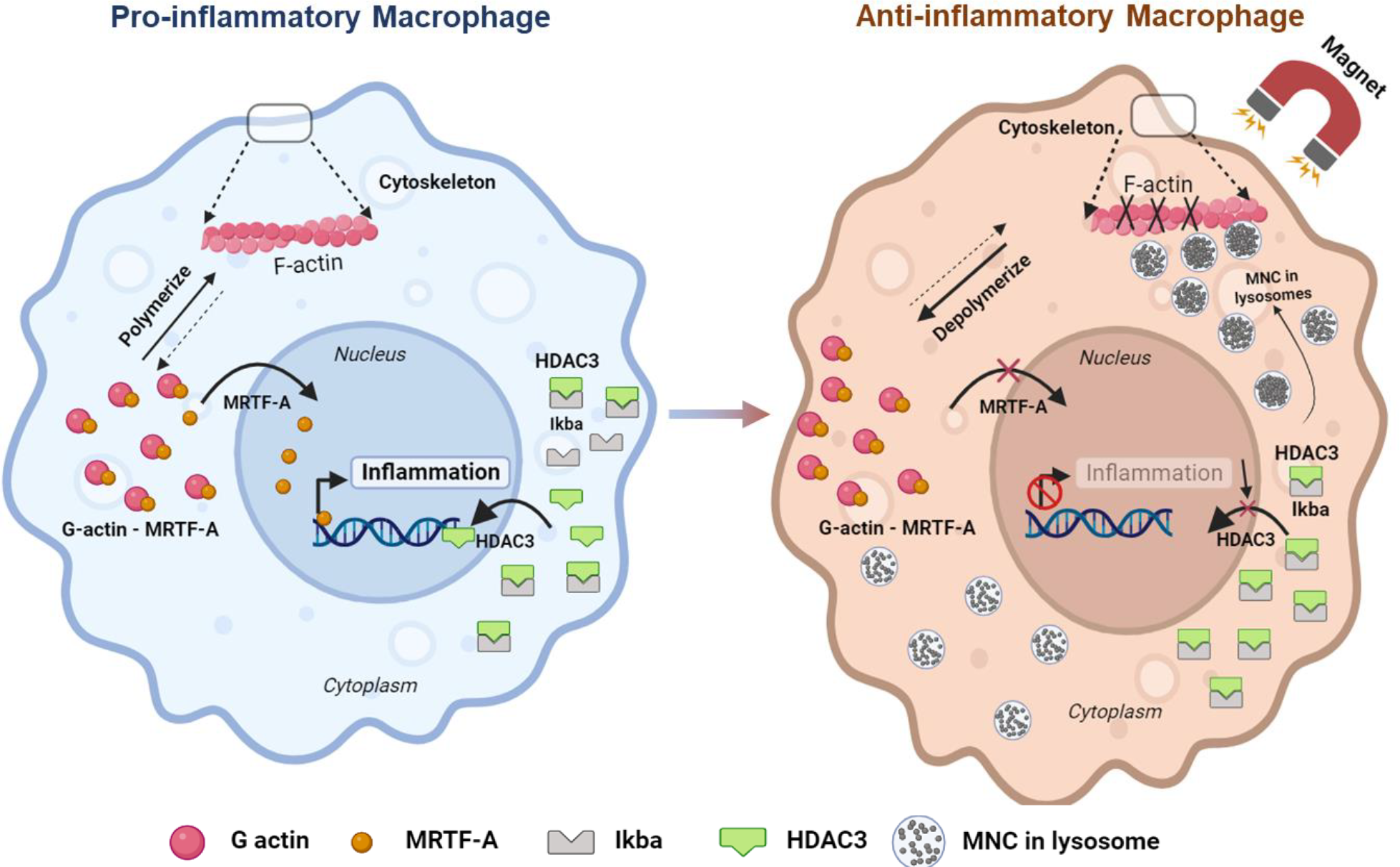
Schematic overview of how magnetic nanocomplexes (MNC) mediated intracellular magnetic force mediates modulation of macrophage phenotype. LPS/IFNү stimulation increases cell spreading of macrophages, during which monomeric G-actin dissociates from MRTF-A to form polymerized F actin, and MRTF-A is translocated to the nucleus. In response to actin polymerization, Ikβα gets degraded, and the unbound HDAC3 is redistributed to the nucleus. MRTF-A and HDAC3 in the nucleus drive major inflammatory gene expression. Pro-inflammatory macrophages readily internalize MNCs and are stored inside the lysosomes. When subjected to the external magnetic field, MNCs are pulled in the direction of the high magnetic field gradient, inducing an intracellular magnetic force on the cytoskeleton. As a result, actin polymerization is disrupted and the nuclear translocation of MRTF-A and HDAC3 are inhibited, thereby limiting inflammatory response and promoting a anti-inflammatory macrophage phenotype.

Overall, our work provided crucial insights into the mechanobiology of macrophages and yielded a potent immunomodulatory therapy for chronic and recalcitrant inflammatory conditions. MNC are clinically translatable materials with FDA-approved formulations available for clinical diagnostic imaging of tumors, cell tracking, and thermal therapy for cancer^31^. Our intracellular approach of applying intracellular magnetic force is unique and is applicable for various wound healing scenarios. This strategy is a paradigm shift in wound healing, and we believe it has the potential to address the limitations of current strategies by instigating localized and targeted effects on immune cells.

## 2. Materials and Methods

### 2.1. Fabrication and characterization of magnetic nanocomplexes (MNC)

MNC are fabricated in three consecutive steps: magnetite nanocrystal synthesis, polymer coating, and functionalization. First, magnetite nanocrystals of the desired size were synthesized by thermo-decomposition of iron acetylacetone^41, 42^. The size of the nanocrystals was confirmed through Transmission Electron Microscope (TEM, FEI Talos F200X). Then, to make them monodisperse in water, nanocrystals are coated with a mixture of phospholipids-polyethylene glycol copolymer (DSPE-PEG2K, Avanti Lipids) and DSPE-PEG2K-maleimide using a dual-solvent exchange method (DSE)^43^. MNC were also labeled with a fluorophore DiI or DiR (Invitrogen) for fluorescent tracking by mixing them in deionized water at a weight ratio of 20:1 for 30 min. The unbound fluorophores were removed by passing the solution through a 0.2 um syringe filter with HT Tuffryn membrane (Pall Inc.). The hydrodynamic size distribution and the zeta potential of the MNC were analyzed using Nanosight 300 (Malvern Analytical) and ZetaView® BASIC NTA (Particle Metrix). The fluorescence intensity of MNC standards with defined iron concentration were measured using a spectrofluorometer for *in vitro* and *in vivo* quantification purposes. Transmission electron microscope (TEM) was used to characterize the size distribution of magnetite nanocrystal and morphology of cells containing MNC. The size of the nanocrystals was then calculated by processing the images using ImageJ software (NIH).

### 2.2. MNC magnetization and force simulation

The saturation magnetization (m_v_) of the nanocrystals measured using a superconducting quantum interference device (SQUID) showed 117.3 emu/g Fe at room temperature^44^. To numerically calculate the magnetic force on MNC, each MNC is approximated to a point dipole. A MATLAB code was written to simulate the magnetic field and magnetic force generated by 1/10” NdFeB rare-earth magnets cylindrical magnet (*B*_*0*_ = 1.3 T) on a single nanocrystal and cells containing specific concentrations of MNC. For calculating the magnetic force on MNC linear superposition principle was applied, and each MNC is considered as a point dipole *M* whose magnetic field is given by:

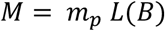

where *m*_*p*_ is the mass of the nanocrystal determine using TEM images (*m*_*p*_ *= V*_*p*_ *ρ, ρ* -density of magnetite = 5.17 x10^3^ g/cm^3^) and *L(B)* is the Langevin function of B, the magnetic field density of a block magnet calculated by integrating over the volume:

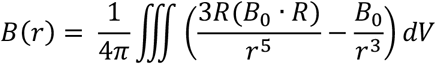

where ***R*** is the vector from the dipole to the point, *r* is the distance, and *B*_*0*_ is the residual magnetization of the magnet. The magnetic force *F*_*m*_ exerted on MNC is calculated through the equation:

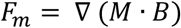

### 2.3. Cell culture, polarization, and effect of intracellular magnetic force

A commercially available mouse peritoneal macrophage cell line (IC-21, ATCC) with C57BL/6 background was used for all *in vitro* studies. These cells are similar to normal mouse macrophages, express macrophage-specific antigens, and are functionally active as normal primary peritoneal macrophages^45^. The macrophages were cultured in RPMI 1640 Medium (ATCC) growth media supplemented with 5% fetal bovine serum (FBS, ThermoFisher) and 1% Antibiotic -Antimycotic (Gibco). For characterizing MNC uptake and force modulation, macrophages were cultured in 35 mm diameter glass bottom (0.17 mm) dishes (Ibidi) until 60-70% confluency. To create proinflammatory (M1-like) macrophage phenotypes, the culture media was supplemented with interferon-gamma (IFNγ, 50 ng/mL) and lipopolysaccharide (LPS, 100 ng/mL). Likewise, to create anti-inflammatory/prohealing (M2-like) macrophages, media was supplemented with interleukin-4 (IL-4, 40 ng/mL) and Interleukin-13 (IL-13, 20 ng/mL). Unstimulated macrophages (M0) cultured in growth media served as controls. Cells were polarized for at least 24 hours before all studies. To investigate the effect of the magnetic force on macrophage culture, two different assemblies of NdFeB rare-earth magnets (K&J magnetics) were used to generate force fields beneath the cells. The cells were cultured in a 0.17 mm thick glass-bottom dish (35 mm diameter, Ibidi) or microflow chamber slide (5 mm width, Ibidi) and placed on top of an array of cylindrical (1/10” dia x 2/10”, *B*_*res*_ = 1.3 T) or rectangular magnets (3/4” x 1/8” x 1/8”, *B*_*res*_ = 1.3 T) respectively. This setup was kept undisturbed in the CO_2_ incubator, and samples were collected at different time points for further analysis.

### 2.4. PrestoBlue assay

The biocompatibility of MNC was investigated by quantifying metabolic performance of macrophages cultured with sterile filtered (0.2 µm) MNC suspension (5-100 µg/mL) in macrophage growth media. The metabolic performance of macrophages was measured using PrestoBlue™ reagent (Invitrogen). Briefly, cells were seeded in a 24-well plate until 60-70% confluency and treated with MNC (5-100 µg/mL). At specific time points, PrestoBlue™ reagent was directly added to the culture wells, incubated for 30 min, and fluorescence was measured at Ex/Em 560/590 nm. Relative fluorescence was compared to the untreated control to assess the effect of MNC.

### 2.5. Ferrozine assay

Ferrozine assay was used, as described previously^46^, to determine the total number of MNC internalized by macrophages over time. Briefly, macrophages were polarized to M0, M1, or M2 phenotypes as described above and incubated with MNC at specific concentrations for 2-24 hours. The cells were then washed with warm PBS, detached using TrypLE™ (Gibco) or scraping in ice, centrifuged at 1,000 g for 10 min, the supernatant discarded, and dehydrated overnight in a vacuum chamber. Then, 50 µl of 12 M HCl was added to the dried pellet followed by 56 µl of 10 M sodium hydroxide (Sigma), 100 µl of 4 M ammonium acetate (Sigma), and 100 µl of hydroxylamine hydrochloride (Sigma). The final volume was made up to 0.5 ml by adding distilled water. After 30 minutes of incubation, the solution was mixed with 0.02% ferrozine solution (Sigma). The absorption of the solution was read at 562 nm with 810 nm as reference. Standards of iron (III)-oxide solutions were used to plot the calibration curve. The iron content in each MNC was calculated using the size of the nanocrystals determined in TEM and density of 5.17 g/cm^3^ with the assumption that the nanocrystals consisted of pure magnetite.

### 2.6. Griesse assay

To determine whether MNC induced generation of nitric oxide in proinflammatory macrophages, the Giresse assay was performed. Macrophages were first cultured in 24 well plates and polarized to proinflammatory phenotype using LPS/IFN treatment. After 24 hours of polarization, the cells were washed twice with 1X PBS and then supplemented with sterile filtered MNC (50 µg/mL) in macrophage growth media for 6 hours. Then, the MNC containing media was replaced with fresh culture media and placed in the humidified incubator. After 24 hours, the macrophage culture supernatants (150 µL/well) were transferred to a fresh 96-well flat-bottom plates. The Griess assay was performed as per manufacturer instruction (Griess Reagent kit, #G7921). Briefly, an equal volume of N-(1-naphthyl) ethylenediamine (Component A) and sulfamic acid (Component B) was first mixed to form the Griess Reagent. The Griess reagent (20 µL/well) was then added to the wells containing culture media, incubated for 30 mins and then the absorbance was measured at 540 nm. NaNO_2_ was used for deriving the standard curve and the results are presented in mean micromolar concentration of nitric oxide in triplicate cultures.

### 2.7. Drug treatment and mechanistic studies

Drug treatment studies were conducted to selectively perturb or enhance pathways to validate the effect of MNC-mediated cytoskeletal modulation. Actin depolymerization study was carried out by supplementing culture media with 100 nM Latrunculin-A (Sigma) for 1 hour, while HDAC3 inhibition studies were carried out by supplementing 20 µM RGFP966 (Sigma) for 24 hours before adding M1 polarizing media (LPS+IFNγ) containing RGFP966. After 24 hours of polarization, cells were collected for RNA extraction and gene expression.

### 2.8. Flow cytometry

Flow cytometric analysis of macrophages was done to analyze phenotype through the expression of specific markers. First, macrophages stimulated using physical or biochemical treatment were detached by non-enzymatically treating them with ice-cold 10 mM phosphate-buffered saline (PBS, Ca^2+^ and Mg^2+^ free, Gibco) for 20-30 min. Then the cell suspension was collected, centrifuged at 300 g for 3 min, fixed in 10% formalin (Z-fix) for 10 min at room temperature, and washed thrice with 10 mM PBS. Then, the cells were permeabilized by adding 0.05% Triton X-100 in 10 mM PBS for 2 mins, washed twice in 10 mM PBS, and incubated in antibody solution for at least 30 minutes at room temperature. Antibodies for iNOS-PE (0.24 µg/test, Santa Cruz) and Arg1-APC (4 µg/test) in 0.5% BSA in PBS were used to delineate macrophage phenotypes. The samples were analyzed in a BD Facsymphony™ A3 cell analyzer (BD Biosciences), and data are analyzed using FlowJo software.

### 2.9. Gene expression

Expression of inflammation, osteogenesis, vasculogenesis, and ECM gene markers were analyzed using qPCR to investigate MNC treatment compared to biochemically stimulated positive controls and untreated negative controls. Briefly, RNA was isolated from cultured cells using Trizol reagent (Invitrogen) and converted to cDNA using superscript™ III Platinum™ one step qRT-PCR Kit according to the manufacturer’s instruction. qPCR was performed on QuantStudio3 (ThermoFisher) using Taqman probes *Tnf* (Mm00443260_g1), *Il1b (*Mm00434228_m1), *Il6 (*Mm00446190_m1), *Nos2* (Mm00440502_m1), *IL-12b (*Mm00434174_m1), *IL-12b (*Mm00434174_m1), *Ccl2 (*Mm00441242_m1), *Cxcl2 (*Mm00436450_m1), *Tlr4 (*Mm00445273_m1), *Retnla (*Mm00445109_m1), *Arg1 (*Mm00475988_m1), *Mrc1(*Mm00485148_m1), *Chil3 (*Mm00657889_mH), *Pparg (*Mm01184322_m1), *Il-10 (*Mm00439616_m1), *Bmp2 (*Mm01340178_m1), *Vegf (*Mm00437306_m1), *Tgfb1 (*Mm01178820_m1), *Osm (*Mm01193966_m1), *Igf1* (Mm00439560_m1) were used for the analysis. *Gapdh (*Mm99999915_g1) was used as internal control.

### 2.10. Immunofluorescence

For immunofluorescence (IF) staining of *in vitro* cultures, the samples were fixed in 10% formalin (Z-fix), washed thrice with 1X PBS, permeabilized with 0.05% Triton X-100 in 10 mM PBS (Sigma) for 2 min, blocked with 1% bovine serum albumin (BSA) in 10 mM PBS for 1 hour and stained with fluorescence antibodies. F actin and G actin of the cells were stained using phalloidin-AF488 (1:400 dilution, ThermoFisher), DNAse I AF594 (9 µg/mL, ThermoFisher) in 1% BSA in PBS respectively for 1 hour. Arginase1 and iNOS are stained using the fluorophore-conjugated antibodies iNOS AF-488 (1:100 dilution, ThermoFisher) and Arginase1(1:50 dilutions, ThermoFisher) in 1% BSA in PBS. Other antibody stains include MRTF-A AF-488 (1:300 dilution, SCBT) and HDAC3 AF-594 (1:300 dilution, SCBT). DAPI (1:500 dilution) was used as a nuclear counterstain. Stained samples were then imaged using an inverted confocal microscope (Nikon A1R).

### 2.11. Migration assay

To make fibrin hydrogel, fibrinogen stock (4 mg of clottable protein/mL of BMEM, bovine plasma, Type I-S, Sigma) was filter sterilized and mixed with 0.5 U/mL thrombin. The mixture was quickly cast in a 35 mm culture dish to form a thin fibrin gel and placed in a humidified incubator for 30 mins to allow complete gelation. The fibrin hydrogel was washed twice with 1X PBS and seeded with MNC internalized macrophages and incubated overnight in humidified incubator for cell attachment. After overnight incubation, MNC internalized macrophages seeded on fibrin hydrogel were placed on an array of cylindrical magnets for 24 hours to exert force on the MNC and the control was the MNC containing macrophages seeded on fibrin hydrogel not subjected to an external magnetic field. The longitudinal movement of macrophages through the fibrin hydrogel due to external magnetic field was measured by capturing Z stack confocal images and the 3D image was constructed using Imaris software.

### 2.12. Micropatterned substrate

The molds for the PDMS stamps were made using a stereolithography-based 3D printer (Form 3, Formlabs). Following printing, the mold was washed in an isopropyl alcohol (IPA) bath (FormWash, Formlabs) for 10 mins and then allowed to air dry at room temperature for 30 mins. The mold was placed in a UV chamber (FormCure, Formlabs) for 30 mins at 60°C temperature. The PDMS stamps were made at a 10:1 ratio of polydimethylsiloxane base to curing agent (Sylgard 184, Dow Corning) and degassed under vacuum for 1 h. The degassed mixture was poured into the 3D printed mold and was incubated at 65°C overnight. The cured PDMS stamp was peeled from the mold, washed with isopropyl alcohol, and treated with air-plasma under vacuum in a plasma cleaner (Harrick Plasma) for making the stamp surface hydrophilic. The stamp was inked with 40 ug/ml fibronectin solution (F0859, Sigma) under sterile condition for 10 mins. Fibronectin stamps were printed onto the surface of hydrophobic cell culture dish (81151, Ibidi) for 5 mins and let the print dry for 10 mins. The surface was then treated with 1% Pluronic F-127 for 10 mins to passivate non-fibronectin-coated regions.

### 2.13. Statistics

All measurements were performed at least in triplicate. Data are plotted as means with error bars representing the standard deviation. The Pearson correlation coefficient (r) was used to evaluate the linear correlation between two variables. Statistical comparisons were made using Student’s t-test (two-tailed and unequal variance) with a 95% confidence limit. Differences with p < 0.05 were considered statistically significant.

## 3. Results

### 3.1. MNC synthesis, fabrication, and characterization

We synthesized magnetite (Fe_3_O_4_) nanocrystals (**Fig.2A**) with a narrow size range of 15 ± 2 nm (**Fig.2B**) through thermo decomposition ^47^. Then we coated the nanocrystal with an amphiphilic phospholipid-PEG copolymer (DSPE-PEG2K) using a dual solvent exchange process to fabricate magnetic nanocomplexes (MNC, **Fig.2C**). The polymer coating onto the hydrophobic nanocrystals renders them water-dispersible while avoiding aggregation. The hydrodynamic diameter of the MNC was found to be 40 ± 3 nm (**Fig.2D)**. To visualize MNC in in-vitro, the MNCs were labeled with fluorescent lipophilic dye DiI (ex/em=549/565 nm) and DiR (ex/em = 710/780 nm), respectively **(Fig.2E)**.

**Figure 2:**
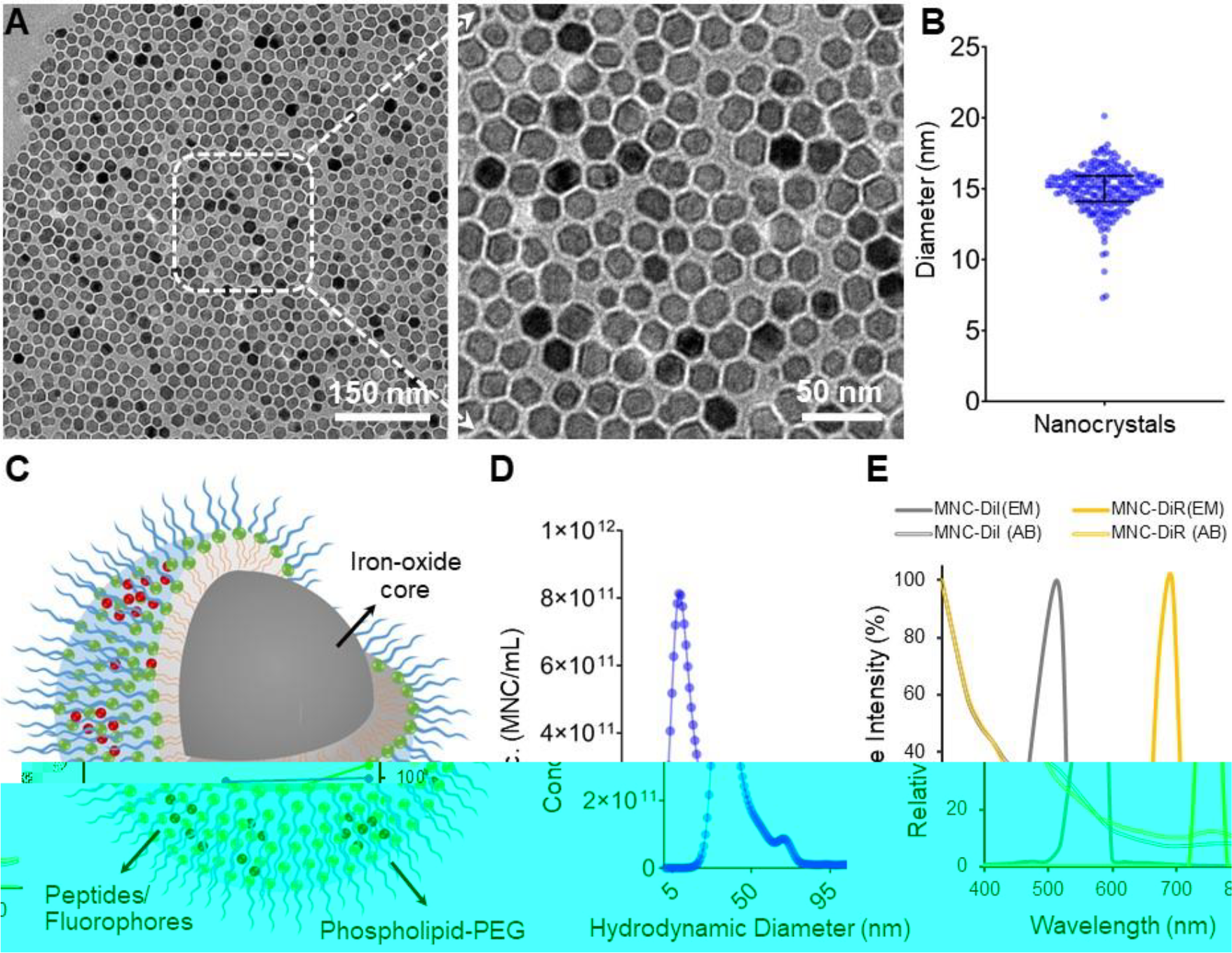
MNC design, fabrication, and characterization. **A)** TEM images of magnetite nanocrystals with uniform **B)** size distribution of 15 nm diameter. **C)** A schematic diagram of magnetic nanocomplexes (MNC) design. MNC was fabricated by coating magnetite nanocrystals with phospholipid-PEG and labeled with lipophilic fluorophores DiI or DiR. **D**) The hydrodynamic diameter of fabricated MNC was measured to be 50 nm diameter. **E)** Absorption and emission spectra of MNCs labeled with DiI and DiR.

### 3.2. MNCs are biocompatible and their uptake rate is phenotype dependent

To assess the biocompatibility of MNC, we first incubated macrophages with linearly increasing concentration of MNCs (0, 2.5, 10, 25, 50, 100 ug/mL) in cell culture media. We performed PrestoBlue assay on Day 1, Day 3 and Day 7 to measure metabolic activity and evaluate cell viability. MNC did not affect the metabolic activity of macrophages at a high concentration of 100 ug/mL at all three timepoints, confirming their cytocompatibility **(Fig.3A)**. To determine the update kinetics of MNC by macrophages, we quantified intracellular MNC at varying their incubation time **(Fig.3B)**. We observed that 80% of the cells were labelled with MNC within 4 hours of incubation and later reached 100% by 12 hours from the flow cytometry analysis **(Fig.3C)**. This indicates that the uptake rate of MNC by macrophage is time dependent. Next, to evaluate whether the phenotype of macrophage influences internalization of MNC, we first treated macrophages with cytokine cocktails LPS/IFN-γ and Il-4/Il-13 to stimulate pro-inflammatory (M1-like) and anti-inflammatory (M2-like) phenotype, respectively. MNCs were added to macrophage cultures (M1 and M2-like) and incubated for 24 hours for uniform uptake. The flow cytometry analysis showed that M1-like macrophages up took significantly more MNCs compared to M2-like, indicating that the internalization of MNC is phenotype dependent **(Fig.3D)**. Next, we verified the MNC uptake pathway by immunofluorescent imaging. The end point of any particle internalized via endocytosis or phagocytosis is lysosome, hence we stained lysosome to determine subcellular localization of MNC. MNCs were colocalized within lysosome around the perinuclear space and were not adhered to the cell surface or trapped in the cell membrane **(Fig.3E)**. The lysosome staining of the macrophage without MNC was used as control (**Supplementary Fig.1)**. Furthermore, we investigated whether MNC enhanced inflammatory response by inducing nitric oxide synthesis in LPS/IFNү treated macrophages by performing Griesse assay as nitric oxide is a marker for proinflammatory phenotype. Interestingly, the nitrite levels in macrophages containing MNC were reduced compared to the macrophages without MNC, however the difference was marginal **(Fig.3F**). In the next section, the MNC internalized macrophages were subjected to external magnetic field to investigate the whether the intracellular magnetic force induced measurable impact on macrophage behavior.

**Figure 3:**
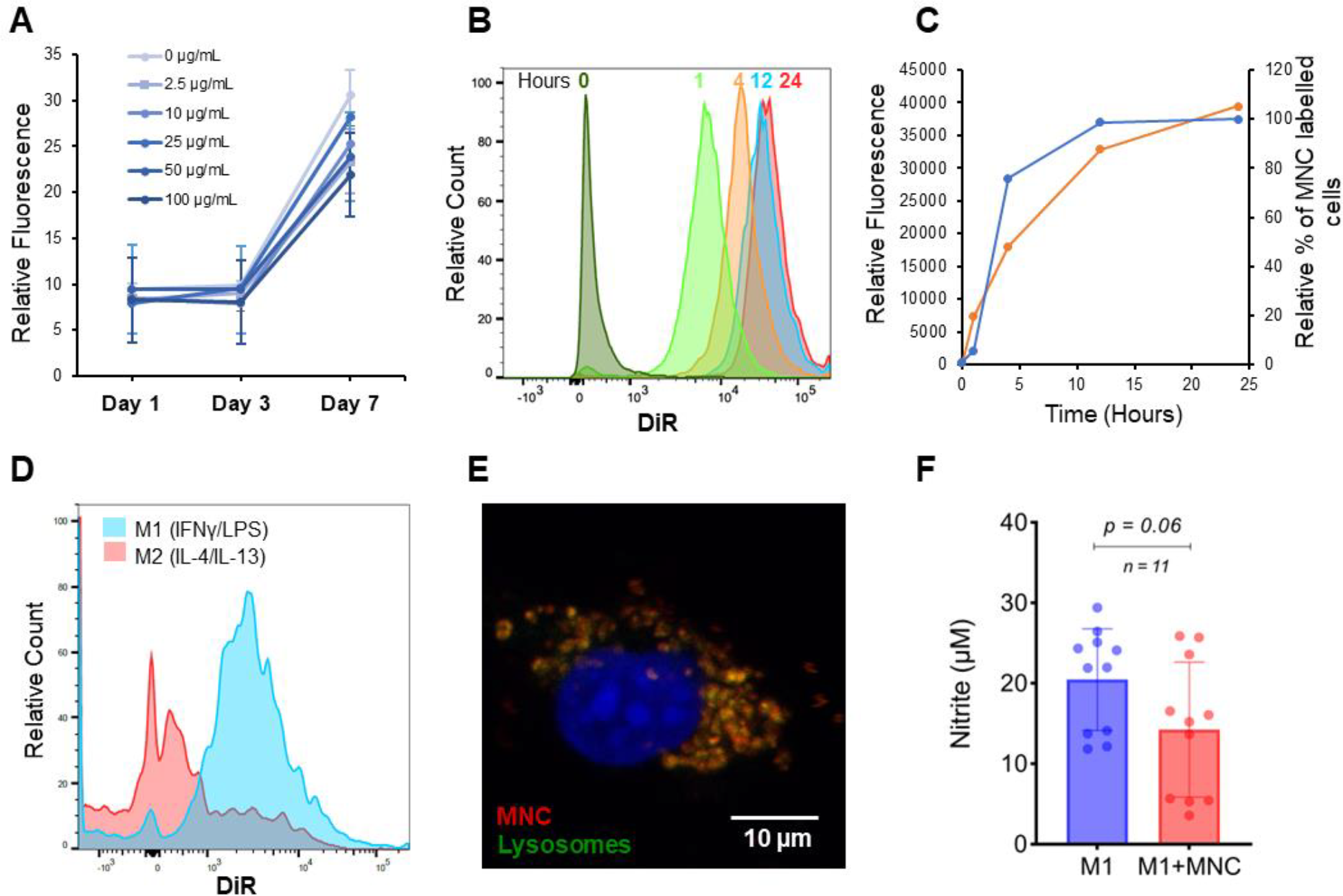
MNCs are biocompatible, and their uptake is phenotype dependent. **A)** Metabolic activity of macrophages incubated with varying concentrations of MNC at days 1, 3, and 7. **B)** Quantification of intracellular MNC at varying incubation periods of 1, 4, 12, and 24 hours. **C)** Plot shows the intracellular fluorescence in the left axis and the percentage of cells internalized MNC in the right axis with respect to time. **D)** Intracellular MNC quantification of M1 and M2-like macrophages **E)** Confocal image shows subcellular localization of MNC in lysosome **F)** Nitrite levels in M1-like macrophage incubated with MNC for 6 hours and control without MNC.

### 3.3. MNC when coupled with an external magnetic field induces intracellular magnetic force

To create intracellular magnetic force in MNC internalized macrophages, we used an array of cylindrical NdFeB magnets as an external magnetic field. The spatial distribution of magnetic field across the surface of a cylindrical magnet was calculated by numerical simulation. The magnetic field gradient was high at the edges of the cylindrical magnets and gradually reduced towards the center. Hence an array of magnets was used to augment magnetic flux distribution. and supply high magnetic field gradient in the axial direction. Assuming a magnetic field gradient of 100 Tm^-1^ the magnetic force acting on an individual MNC is on the order of 0.6 fN and which is ∼ 450-fold higher than the gravitational force **(Fig.4A)**. The amount of MNC internalized by each cell was calculated to be around 3.5×10^6^ based on the iron content measurements obtained from ferrozine assay. The intracellular magnetic force along the direction of the external magnetic field lines experienced by a macrophage containing 3.5 million MNC when placed at the edge of the magnet is calculated as 2.275 nN **(Fig 4B)**.

**Figure 4:**
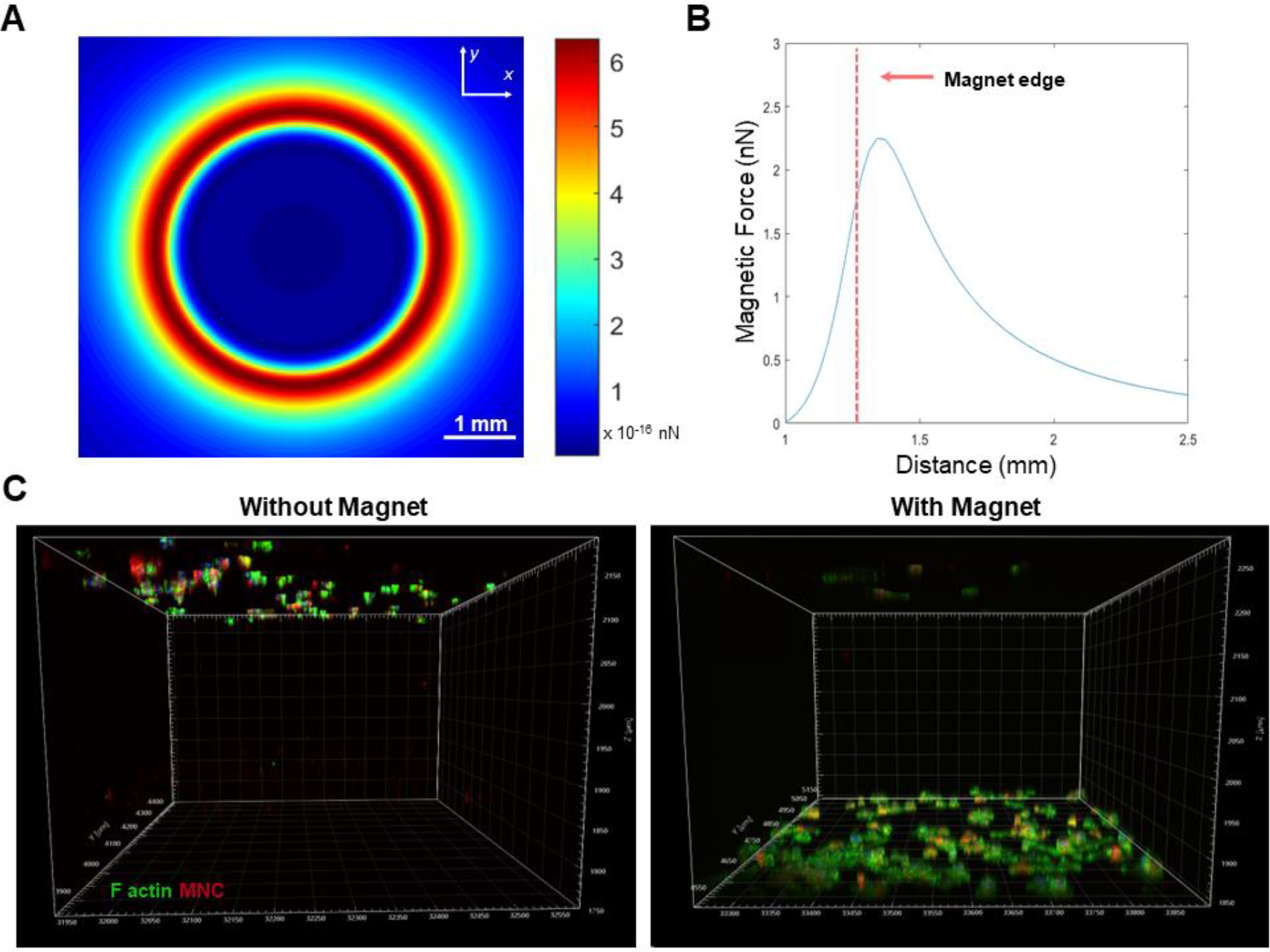
Numerical simulation of magnetic field gradient and intracellular magnetic force. **A)** The magnitude of the magnetic field created by a single cylindrical magnet computed from Maxwell’s equations. **B)** Magnetic force exerted on a single macrophage containing 3.5 million MNC across the cylindrical magnet. **C)** Macrophages containing MNC (red) were stained for F actin (green) did not migrate to the bottom of the culture dish without an external magnet (bottom left). With an external magnetic field, the MNC-containing cells migrated and attached to the bottom (Botton right).

To check the intensity of intracellular magnetic force in response to external magnetic field, we performed a migration assay using fibrin hydrogel. The MNC internalized macrophages were seeded on a thin fibrin hydrogel of 0.5 mm thickness. The MNC loaded macrophages migrated through the fibrin hydrogel and arrived at bottom surface of the culture dish when placed on an array of cylindrical magnets for 24 hours **(Fig 4C. bottom right)**. Whereas, without external magnetic field, the MNC loaded macrophages did not migrate and remained attached to the top surface of the fibrin hydrogel **(Fig 4C, bottom left)** even after 48 hours of incubation. This confirms the ability to specifically target MNC internalized macrophage with biophysical cues using external magnetic field. The MNC internalized macrophages attached to the surface and quantification graph of overall F actin and MNC across the depth of fibrin hydrogel with and without magnet are shown in **Supplementary Fig 2**. We next characterize the correlative changes in actin cytoskeletal dynamics, nuclear to cytoplasm levels of MRTF-A and HDAC3, protein (INOS & ARG1) and differential gene expression in macrophage phenotypes.

### 3.4. Characterization of macrophage immunotypes

To understand the cytoskeleton dynamics and associated functional phenotypes of macrophages, we polarized macrophages to pro-inflammatory (M1-like) and anti-inflammatory (M2-like) phenotype by treating with LPS/IFN-γ and Il-4/Il-13 for 24 hours, respectively. We fixed the cells and stained for F actin and G actin **(Fig 5A)**. We first analyzed cell morphology by measuring cell area and quantified F actin to G actin ratio in unstimulated (M0), proinflammatory (M1-like) and anti-inflammatory (M2-like) macrophage phenotype. The cell area of the M1-like **(Fig 5B)** macrophages was significantly high along with F actin/G actin ratio **(Fig 5C)** compared M0 and M2-like macrophages. When we quantified the nuclear to cytoplasm ratio of actin dependent transcription factor, MRTF-A and chromatin modifying enzyme, HDAC3, M1-like macrophages showed significantly high nuclear levels of MRTF-A **(Fig 5D)** and HDAC3 **(Fig 5E)** compared to M0 and M2-like, correlating to increased F/G actin levels as observed previously. The protein expression profile of the stimulated macrophages was analyzed by staining for prominent M1 markers iNOS and M2 marker ARG1 **(insets in Fig 5A)**. iNOS protein expression levels were significantly high in M1-like, and ARG1 levels were significantly high in M2-like macrophages **(Fig 5 F, G)**. Differential gene expressions of M1 and M2 like macrophages were analyzed by measuring mRNA levels of prominent pro-inflammatory and anti-inflammatory genes by qPCR **(Fig 5H)**. All this experiment together reveals the correlation between actin dynamics and nuclear to cytoplasm redistribution of MRTF-A and HDAC3 and how the correlation is associated with prominent pro-inflammatory and anti-inflammatory protein and gene expression. In the next section, we study the effect of intracellular magnetic force on modulating F/G actin levels and subsequent change in nuclear MRTF-A and HDAC3 levels.

**Figure 5:**
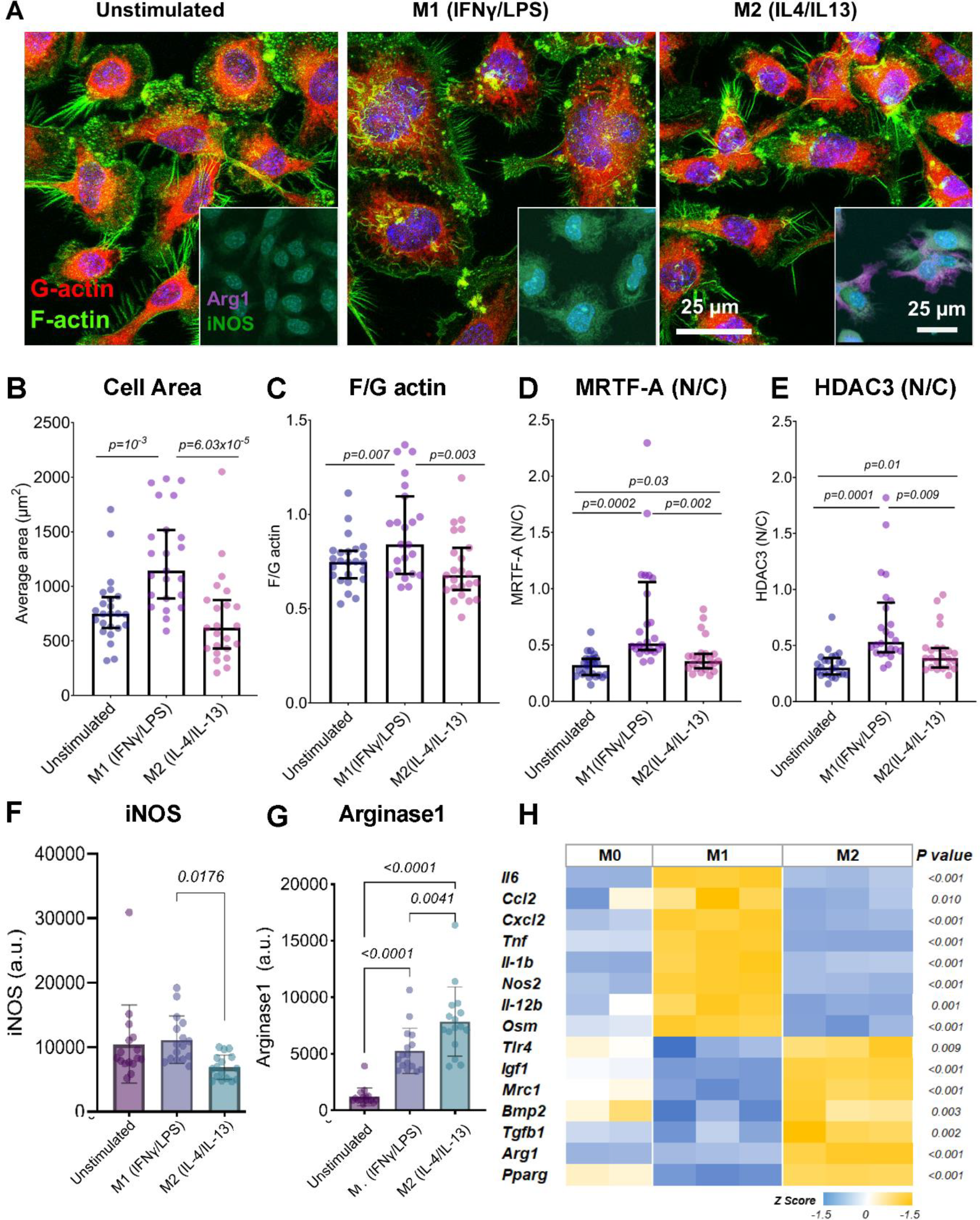
Cytoskeletal, proteomic, and genomic characterization of macrophage phenotypes. **A)** unstimulated, M1-like, and M2–like macrophages stained for F actin (green) and G actin (Red). The insets contain corresponding cells stained for iNOS (green) and Arg1 (magenta). Quantification of **B)** cell area, **C)** F/G actin levels, **D)** nuclear to cytoplasmic levels of MRTF-A and **E)** HDAC3, **F)** iNOS, and **G)** Arginase protein levels in the unstimulated, M1-like and M2-like macrophages. **H)** Heat map representing Z score values of mRNA levels of inflammatory and anti-inflammatory genes in macrophage phenotype. Gapdh was used as an internal control for normalization.

### 3.5. Intracellular magnetic force alters macrophages cytoskeleton by regulating nuclear to cytoplasm redistribution of MRTF-A and HDAC3

Actin cytoskeleton dynamics play a crucial role in cellular processes such as gene transcription, signal transduction and intercellular interaction. To study changes in their dynamic in response to intracellular magnetic force, we added MNCs to the M1-like macrophages and incubated them for 6 hours for internalization. We then placed the MNC internalized macrophages on an array of cylindrical magnets for 24 hours to exert intracellular force. The MNC containing macrophages without an external magnet was assigned as control. The cell shape of macrophages placed on the junctions between the cylindrical magnets appeared slightly elongated due to the pull of intracellular MNC in the direction of external magnetic field gradient lines **(Fig.6A)**.

We quantified the effect of magnetic field gradient on intracellular MNC by measuring the distribution of MNC within the cytoplasm. The intracellular MNC distribution factor was significantly altered with external magnet compared to control without magnet **(Fig.6B)**. We next evaluated how the MNC redistribution within the cytoplasm has altered the geometry of cell by measuring cell circularity index. The exposure to external magnetic field has significantly reduced the circularity index of the macrophages compared to control without magnets. (Circularity Index = 4*Pi*cell area /cell perimeter^2)

We next assess whether the MNC mediated intracellular force altered cell shape by rearranging actin cytoskeletal organization. From quantifying filamentous F actin and monomeric G actin levels, it was clear that the intracellular magnetic force has significantly reduced F actin to G actin ratio by perturbing actin polymerization **(Fig.6D)**. We stained the macrophages for F actin and G actin and quantified their levels by capturing confocal Z stack images **(Fig 6E)**. We next wanted to determine whether the changes in the actin dynamics regulated nuclear to cytoplasmic translocation of MRTF-A and HDAC3. Interestingly, the nuclear to cytoplasmic levels of MRTF-A **(Fig 6F)** and HDAC3 **(Fig.6G)** were significantly reduced in MNC internalized macrophages with external magnet compared to control. This confirms that the intracellular magnetic force restricts nuclear translocation of MRTF-A and HDAC3 by regulating actin cytoskeletal dynamics. Because MRTF-A and HDAC3 mediates pro-inflammatory transcription, we next checked whether sequestration of MRTF-A and HDAC3 in cytoplasm promoted anti-inflammatory phenotype. We confirmed that the intracellular magnetic force indeed increased ARG1 levels compared to control by performing flow cytometry **(Fig.6H)**. This suggests that MNC induced intracellular magnetic force can downsize actin regulated inflammatory transcription and promote anti-inflammatory phenotype.

**Figure 6:**
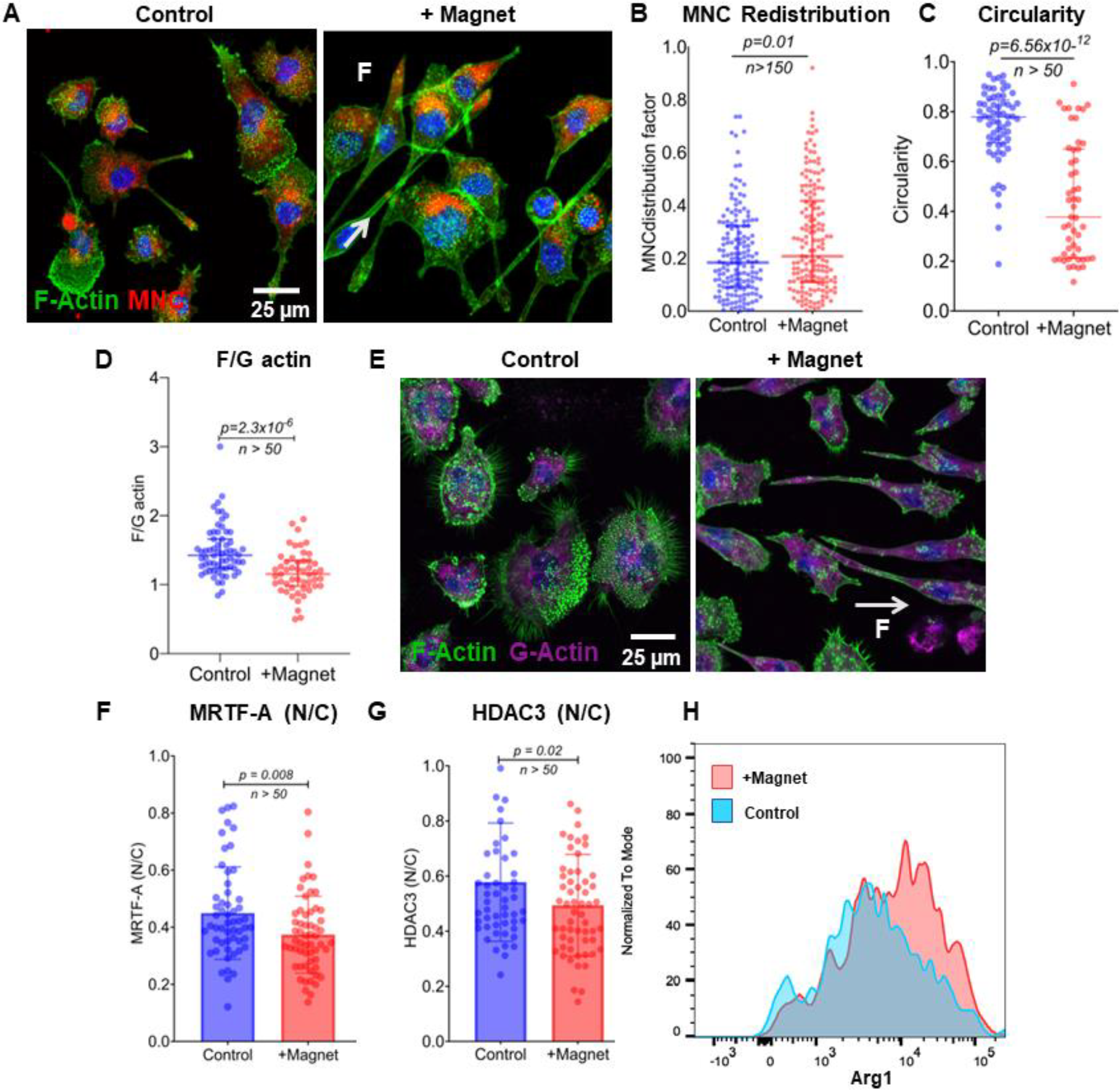
Intracellular magnetic force altered macrophage cytoskeleton dynamics and protein expression. **A)** Confocal images of MNC-loaded macrophages with and without magnet. F actin in green and MNC in red. Intracellular MNCs were pulled along the direction of the external magnetic field gradient. **B)** The intracellular MNC distribution within the cytoplasm was measured from confocal images. **C)** The change in cell shape was quantified by measuring circularity, **D)** A Plot showing F actin and G actin levels between control and with magnet measured from confocal images, **E)** MNC internalized macrophages-stained F actin in green and G actin in magenta. Nuclear to cytoplasmic levels of **F)** MRTF-A and **G)** HDAC3 quantified by immunofluorescent staining **H)** ARG1 expression

Since a strong correlation was observed between actin polymerization and nuclear to cytoplasm levels of MRTF-A and HDAC3, we wanted to gain mechanistic understanding of how inflammatory response is associated with morphological changes and MRTF-A and HDAC3 levels. First, we treated M1-like macrophages with Latrunculin-A (Lat-A) to inhibit actin polymerization and analyzed their effect on cell shape and gene expression. The F actin level was significantly reduced **(Supplementary Fig.3B)**, and the overall MRTF-A levels were significantly high **(Supplementary Fig.3C)**, however there was no significant difference in the nuclear to cytoplasmic levels compared to control M1-like macrophages without Lat-A treatment. Interestingly gene expression analysis showed there was a significant increase in *Bmp2* expression, a prominent osteogenic marker. **(Supplementary Fig.3D)**

We next wanted to examine cytoskeletal changes and gene expression of M1-like macrophages upon HDAC3 inhibition using RGFP966. We treated the M1-like macrophages with RGFP966 and noticed significant decrease in F actin levels **(Supplementary Fig.4B)**, and circularity index **(Supplementary Fig.4C)** upon inhibiting HDAC3, compared to M1-like macrophages with no treatment. Interestingly, along with *Bmp2*, some of the anti-inflammatory genes such as *Chil3, Pparγ* were significantly upregulated when HDAC3 was inhibited **(Supplementary Fig.4D)**. Additionally, we noticed inhibition of actin polymerization significantly reduced iNOS levels **(Supplementary Fig.5B)** and inhibition of HDAC3 significantly increased ARG1 levels **(Supplementary Fig.5C)** compared to control M1-like macrophages with no treatment. Moreover, HDAC3 inhibition significantly increased iNOS to Arg1 ratio **(Supplementary Fig.5D)** promoting anti-inflammatory phenotype.

Taken together, these data show that MNC mediated intracellular magnetic force modifies actin cytoskeletal dynamics to promote a pro-healing macrophage phenotype by facilitating epigenetic alteration (HDAC3 Levels) and regulating transcription factors (MRTF-A). In the next sections, intracellular magnetic force induced changes in gene expression are analyzed using an engineered micropatterned substrate.

### 3.6. MNC mediated transcriptional reprogramming downsized proinflammatory phenotype

As mentioned previously, the magnetic field gradient is not uniform across the surface of external magnet (an array of cylindrical magnets). To determine the regions on the culture surface which experiences high magnetic field gradient when placed on an array of cylindrical magnets, we dispersed MNC in PBS, placed it on an array of cylindrical magnets and waited for a few mins for the MNC to arrange themselves in response to external magnetic field lines. The MNCs gradually accumulated at the junctions between two cylindrical magnets where magnetic field gradient is high **(Fig.7A inset)**. Likewise, the intracellular MNC arranges themselves in the same pattern as shown in **Fig.7A** where M1-like macrophages are stained for F actin in green and MNCs in red.

**Figure 7:**
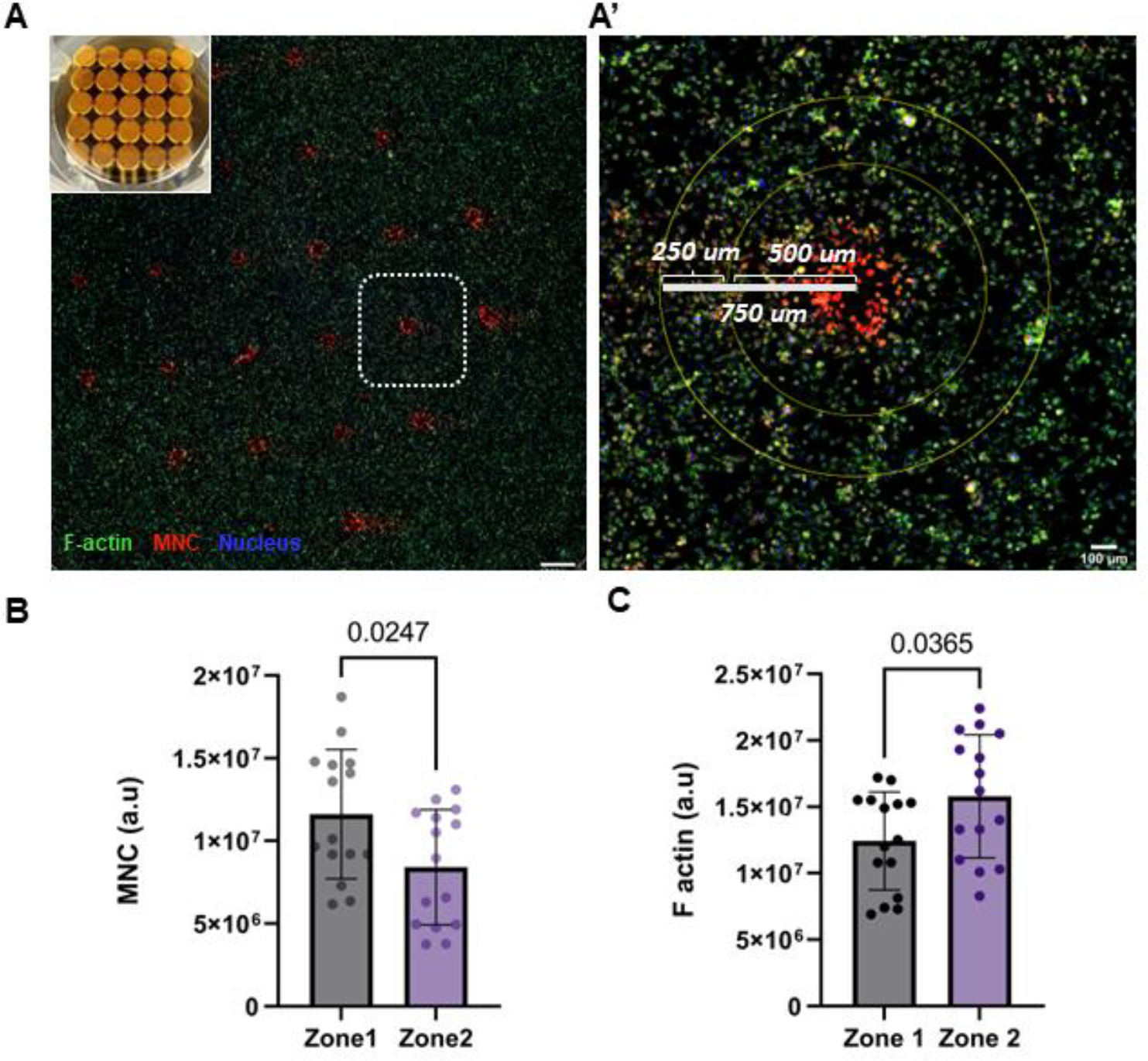
Intracellular MNC experiences a gradient of magnetic field. **A)** M1-like macrophages loaded with MNC (red) when subjected to an external magnetic field, the intracellular MNC accumulated towards the regions of the high magnetic field gradient. **A’)** Enlarged image of dotted box in fig A, which shows the region comprising zone 1 and 2. MNC dispersed in PBS accumulated between the junction of an array of cylindrical magnets (inset). Plots **B)** and **C)** shows the quantifying of intracellular MNC and F actin in zone 1 and 2, respectively.

We defined two zones around the junctions of cylindrical magnet, zone 1 comprising of an inner circle with radii-500 um, zone 2 defined as the region between inner and the outer circle of radii 750 um **(Fig.7A’)**. We measured and compared F actin levels to the amount of intracellular MNC between the two zones. F actin levels were significantly reduced in zone 1 where cells experience maximum magnetic field compared to zone 2, where cells are slightly farther from high magnetic field gradient **(Fig.7B)**. The amount of intracellular MNC was significantly high in cells present in zone 1 compared to zone 2 **(Fig.7C)**. Therefore, we wanted to analyze the transcriptional changes in specific population of cells around these junctions, where magnetic fields gradient is high, to understand the intracellular magnetic force mediated transcriptional reprogramming more accurately. We developed a micropatterned substrate using fibronectin stamps with a geometry that allows cells to adhere and proliferate only in the region subjected to high magnetic fields. Pluronic F127 was used to passivate non-fibronectin printed culture surfaces and inhibited cell attachment **(Supplementary Fig 6A&B)**. Macrophages were seeded and polarized to M1-like phenotype and supplemented with MNC in the micropatterned substrate. When subjected to an external magnetic field, the intracellular MNCs were pulled towards the junction of cylindrical magnet whereas in the control without magnet, macrophages with intracellular MNCs were evenly distributed **(Fig 8A)**. The gene expression analysis showed that *Nos2*, a prominent pro-inflammatory marker was significantly reduced in MNC internalized M1-like macrophages coupled with magnet compared to control. Interestingly, *Tgf-β*, an anti-inflammatory marker showed an increasing trend *(p=0*.*06)* due to intracellular magnetic force compared to control without external magnetic field **(Fig 8B)**. The gene expression study using unpatterned substrate did not show significant difference between control and with external magnetic field (Fig 8C) for the same set of genes, confirming the importance of the engineered micropatterned substrate for accurate transcriptional analyses. Overall, MNC mediated intracellular force can promote pro-healing macrophage phenotype by harnessing cytoskeleton dependent transcriptional reprogramming.

**Figure 8:**
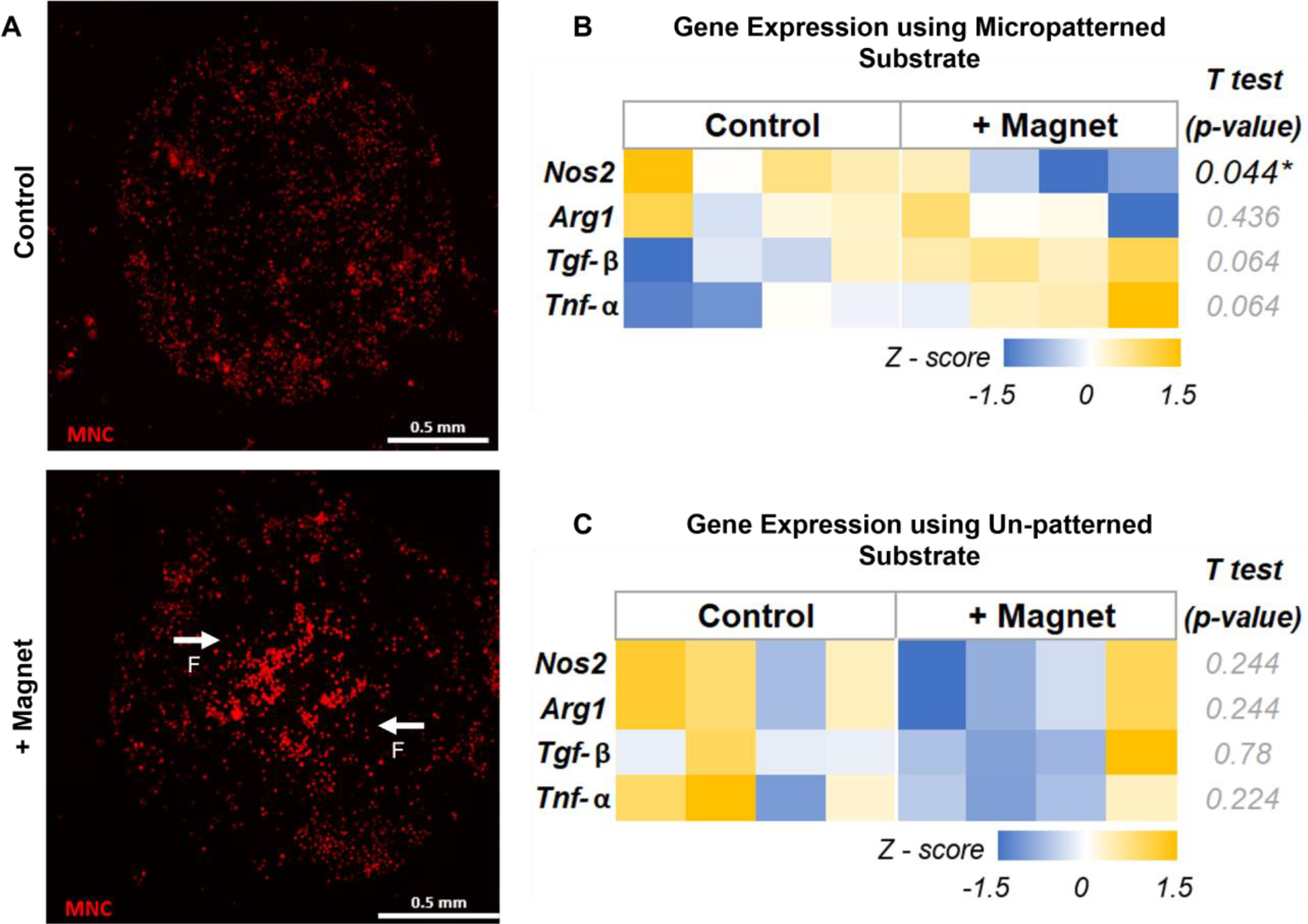
Intracellular magnetic force induces transcriptional reprogramming. **A)** Fluorescent image showing the arrangement of Intracellular MNC on a micropatterned substrate without and with the magnet. **B)** and **C)** Heatmap representing Z score values of mRNA levels of genes without and with the magnet using micropatterned and unpatterned substrates, respectively. Gapdh was used for normalization.

## 4. Discussion

Chronic inflammation is a common pathological basis for various age-related diseases including atherosclerosis, diabetics, cancer, Alzheimer’s and nonunion. Macrophages are the key inflammatory cells, and play a crucial role in tissue development, homeostasis, and repair. Some of the chronic wounds fail to heal despite therapeutic interventions due to delay or improper transition from inflammatory to reparative phase in the healing process. This is attributed either to dysregulated or prolonged expressions of pro-inflammatory macrophage phenotype^48^. Though modulating macrophage phenotype using biochemical cues may temporarily aid the healing, they often result in off target effects or aimed at suppressing single inflammatory cytokines and it is also challenging to control their release kinetics in in-vivo spatially and temporally^49^. Hence there is a need for multi-dimensional approach to modulate macrophage phenotype such that it enhances tissue regeneration and promotes wound healing.

Here we have developed an immunomodulatory strategy using intracellular magnetic force which shifts pro-inflammatory macrophage phenotype towards pro-healing. To our knowledge, we are the first to show that intracellular biophysical cues can be employed to stimulate phenotypic transition without biochemical cues. We have demonstrated that MNCs design and fabrication supports their biocompatibility and the cellular uptake rate of MNC is dependent on macrophage immunotype. MNCs were preferentially internalized by M1-like pro-inflammatory macrophages without affecting cellular metabolism and viability. After internalization, the MNCs were localized in lysosomes around the peri nuclear space. Reactive oxygen species and nitric oxide are synthesized by macrophages during the inflammatory phase in a TLR mediated manner to eliminate intracellular bacteria and fight infections ^50^. The M1-like macrophages that internalized and stored MNC in lysosome did not stimulate nitric oxide synthesis and enhance inflammatory response. On the contrary, the nitric oxide levels were decreased within 24 hours of MNC internalization*(p=0*.*06)*. This supports the cytocompatibility of MNC and by itself MNC do not promote pro-inflammatory macrophage phenotype.

Researchers have shown that disruption force to break actin and its binding proteins such as filamin and alpha actinin to be in the range of 40-80 pN^51^ The force at which an actin filament breaks was measured to be 108 pN, regardless of filament length^52^. From numerical simulation, we calculated the force exerted on 1 million MNC within the cytoplasm to be 0.6 nN and from ferrozine assay, we calculated that each macrophage incorporated around 3.5 million MNCs and exerted an intracellular magnetic force of 2.27 nN. This amount of force was sufficient to pull the MNC loaded macrophages through the fibrin hydrogel when subjected to external magnetic field within 24 hours as well as to disrupt actin polymerization. This finding confirms the possibility of MNC labelled macrophages to be retained and targeted within inflamed tissues *in vivo* studies.

Proinflammatory macrophages (M1 like) containing MNCs when subjected to an external magnetic field altered intracellular MNC redistributed within the cytoplasm and modified macrophage shape to more elongated and significantly less circular compared to control. Several studies have shown that confining macrophages by culturing in a 20 µm wide area, decreased their cell spreading area, resulting in increased ARG1 expression levels^38^. Similarly culturing M1-like macrophages in a 20-30 µm microporous substrate, downsized actin regulated transcription along with associated proinflammatory gene expression. Confinement of the cell spreading area was also shown to limit HDAC3 and MRTF-A derived inflammatory gene expression^28^ Likewise, MNC induced intracellular force significantly decreased F actin to G actin ratio and increased ARG1 expression confirming the influence biophysical cue on macrophage functional phenotypic change. The nuclear to cytoplasmic ratio of MRTF-A and HDAC3 were significantly reduced due to perturbation of actin polymerization by MNC. MRTF-A is bound to G actin in the cytoplasm and upon F actin polymerization, MRTF is cleaved and translocated to nucleus to drive structural and inflammatory gene expression. MRTF-A has been shown to enhance NF-kβ dependent pro-inflammatory transcription in macrophage. MRTF-A deficiency is associated with repressed chromatin structure with enriched acetylated histone H3 and H4. MRTF-A has shown to play vital role in positioning and recruitment of p65 to the nucleus and interact with histone modifying enzymes to drive inflammatory gene expression^53^. Similarly, several studies have reported that HDAC3 promotes inflammatory macrophage phenotype by deacetylating NF-kβ p65 activity ^54^

Mechanistic studies shown here reinforce that selective inhibition of HDAC3 significantly increased ARG1 expression levels and significantly decreased F actin levels. Interesting HDAC3 inhibition altered the cell shape to a more elongated phenotype like IL4/IL-13 treated M2-like macrophages and, increased the expression of *Bmp2* and some anti-inflammatory genes like *Chil3* and *Pparγ*. Inhibition of F actin polymerization also increased *Bmp2* gene expression, but significantly decreased iNOS levels. It is of particular importance to note that our data suggests MNC mediated intracellular magnetic force on actin cytoskeletal regulates nuclear to cytoplasm shuttle of MRTF-A and HDAC3, which subsequently reduces iNOS and increase ARG1 protein levels, to promote pro-healing phenotype of macrophages.

One of the major findings in this report is the significant reduction of *Nos2* mRNA levels in a narrowed down population of MNC internalized M1-like macrophages subjected to high magnetic field using our engineered micropatterned substrate. We speculate that the intracellular magnetic force disrupted F actin polymerization and prevented MRTF-A and HDAC3 translocation into the nucleus. Sequestration of MRTF-A in the cytoplasm stalls the recruit of p65, a subunit of NF-kβ to the nucleus. On the other hand, reduced nuclear to cytoplasm ratio of HDAC3 could have increased *Pparγ* which may inhibit LPS-induced secretion of NF-kβ by upregulating Ikβα, a negative regulator of NF-kβ stimulation^55^. It is central to note that the inhibition of actin polymerization and HDAC3 increased *Bmp2* gene expression levels, which is belongs to Tgf-β superfamily, and promotes bone formation by differentiating MSC to osteoblasts and osteocytes that produce extracellular matrix for bone tissue formation^56^ Our data also implicate that Intracellular magnetic force increases *Tgf-β* mRNA levels *(P=0*.*06)* which may be due to MNC mediated inhibition of HDAC3 nuclear translocation.

In essence, our report indicates that the MRTF-A interaction with NF-kβ is responsible for regulating proinflammatory transcription. However, it is paramount not to overstate this conclusion, because MRTF-A also interacts with other transcription factors such as AP-1 and SRF to drive genes responsible for inflammation. The current study focuses entirely on the F actin and G actin cytoskeletal dynamics. But the key mechanism to generate branched network is dependent on Actin related protein 2/3 complex (Arp2/3 complex) and effect of biophysical cue on Arp2/3 complex are not discussed here. The MNC mediated changes in the cytoskeletal genes involved in actin polymerization and their influence on regulating MRTF-A and HDAC3 nuclear translocation are not discussed here and needs further exploration.

Overall, MNCs are a potent tool to manipulate actin cytoskeleton-driven transcription machinery to modulate inflammation. Biophysical manipulation through engineered MNC can elicit transcriptional control of macrophage phenotype by regulating cytoplasm to nuclear translocation of actin cytoskeleton-dependent transcription factors (MRTF-A) and epigenetic markers (HDAC3), which drives the expression of inflammatory and osteogenic genes, respectively. These findings may facilitate efforts to develop MNC-based resolution-centric therapeutic intervention to direct macrophage phenotype and function towards healing and tissue regeneration, to supplement or replace the currently used anti-inflammatory therapies. Our work provides crucial insights into mechanobiology of macrophages and yields a potent immunomodulatory therapy for nonunion.

## 5. Acknowledgment

Research reported in this publication was supported in part by the National Institute of Arthritis and Musculoskeletal and Skin Diseases (NIAMS) Award Number R21AR078447, National Institute of General Medical Sciences (NIGMS) of the National Institutes of Health under Award Numbers P20GM130456 and P20GM103436-20 (KY IDeA Networks of Biomedical Research Excellence), Small grant award by the UK center for Clinical and Translational Science (CCTS) and Orthopedic Trauma Association (OTA, Grant Number: 6889). The content is solely the responsibility of the authors and does not necessarily represent the official views of the National Institutes of Health or other grant funding agencies. We greatly appreciate Dr. Christine Trinkle and Shaikh Al Mahmud Bhuiyan for their guidance and generous help with 3D printing and developing PDMS stamps. We are deeply grateful to Dr. Ashley Seifert for his guidance and offering lab space to develop and conduct experiments involving micropatterned substrate.

## Reference

1. Barbour, K.E., Helmick, C.G., Boring, M. & Brady, T.J. Vital Signs: Prevalence of Doctor-Diagnosed Arthritis and Arthritis-Attributable Activity Limitation - United States, 2013-2015. MMWR Morb Mortal Wkly Rep 66, 246–253 (2017).

2. Loi, F. et al. Inflammation, fracture and bone repair. Bone 86, 119–130 (2016).

3. Gu, Q., Yang, H. & Shi, Q. Macrophages and bone inflammation. Journal of Orthopaedic Translation 10, 86–93 (2017).

4. Chang, M.K. et al. Osteal Tissue Macrophages Are Intercalated throughout Human and Mouse Bone Lining Tissues and Regulate Osteoblast Function In Vitro and In Vivo. The Journal of Immunology 181, 1232–1244 (2008).

5. Vi, L. et al. Macrophages Promote Osteoblastic Differentiation In Vivo: Implications in Fracture Repair and Bone Homeostasis. Journal of Bone and Mineral Research 30, 1090–1102 (2015).

6. Jilka, R.L., Weinstein, R.S., Parfitt, A.M. & Manolagas, S.C. Perspective: Quantifying Osteoblast and Osteocyte Apoptosis: Challenges and Rewards. Journal of Bone and Mineral Research 22, 1492–1501 (2007).

7. Lee, S.K. & Lorenzo, J. Cytokines regulating osteoclast formation and function. Curr Opin Rheumatol 18, 411–418 (2006).

8. Chan, J.K. et al. Low-dose TNF augments fracture healing in normal and osteoporotic bone by up-regulating the innate immune response. EMBO Molecular Medicine 7, 547–561 (2015).

9. Kon, T. et al. Expression of Osteoprotegerin, Receptor Activator of NF-κB Ligand (Osteoprotegerin Ligand) and Related Proinflammatory Cytokines During Fracture Healing. Journal of Bone and Mineral Research 16, 1004–1014 (2001).

10. Khanna, S. et al. Macrophage Dysfunction Impairs Resolution of Inflammation in the Wounds of Diabetic Mice. PloS one 5, e9539 (2010).

11. Mirza, R. & Koh, T.J. Dysregulation of monocyte/macrophage phenotype in wounds of diabetic mice. Cytokine 56, 256–264 (2011).

12. Hesketh, M., Sahin, K.B., West, Z.E. & Murray, R.Z. Macrophage Phenotypes Regulate Scar Formation and Chronic Wound Healing. Int J Mol Sci 18 (2017).

13. Koh, T.J. & DiPietro, L.A. Inflammation and wound healing: the role of the macrophage. Expert Rev Mol Med 13, e23 (2011).

14. Loi, F. et al. Inflammation, fracture and bone repair. Bone 86, 119–130 (2016).

15. Zerbini, C.A.F. et al. Biologic therapies and bone loss in rheumatoid arthritis. Osteoporosis International 28, 429–446 (2017).

16. Yeo, L. et al. Expression of chemokines CXCL4 and CXCL7 by synovial macrophages defines an early stage of rheumatoid arthritis. Annals of the Rheumatic Diseases 75, 763–771 (2016).

17. Takano, S. et al. Synovial macrophage-derived IL-1βregulates the calcitonin receptor in osteoarthritic mice. Clinical & Experimental Immunology 183, 143–149 (2016).

18. Zhu, W. et al. Ankylosing spondylitis: etiology, pathogenesis, and treatments. Bone Research 7, 22 (2019).

19. Rao, A.J. et al. Revision joint replacement, wear particles, and macrophage polarization. Acta Biomater 8, 2815–2823 (2012).

20. Hamilton, J.A. & Tak, P.P. The dynamics of macrophage lineage populations in inflammatory and autoimmune diseases. Arthritis & Rheumatism 60, 1210–1221 (2009).

21. Talpin, A. et al. Monocyte-derived dendritic cells from HLA-B27+ axial spondyloarthritis (SpA) patients display altered functional capacity and deregulated gene expression. Arthritis research & therapy 16, 417 (2014).

22. Leipe, J. et al. Role of Th17 cells in human autoimmune arthritis. Arthritis & Rheumatism 62, 2876–2885 (2010).

23. Bitar, D. & Parvizi, J. Biological response to prosthetic debris. World J Orthop 6, 172–189 (2015).

24. Schlundt, C. et al. Macrophages in bone fracture healing: Their essential role in endochondral ossification. Bone 106, 78–89 (2018).

25. Zhang, R., Liang, Y. & Wei, S. M2 macrophages are closely associated with accelerated clavicle fracture healing in patients with traumatic brain injury: a retrospective cohort study. Journal of Orthopaedic Surgery and Research 13 (2018).

26. Alhamdi, J.R. et al. Controlled M1-to-M2 transition of aged macrophages by calcium phosphate coatings. Biomaterials 196, 90–99 (2019).

27. Wójciak-Stothard, B., Madeja, Z., Korohoda, W., Curtis, A. & Wilkinson, C. Activation of macrophage-like cells by multiple grooved substrata. Topographical control of cell behaviour. Cell biology international 19, 485–490 (1995).

28. Jain, N. & Vogel, V. Spatial confinement downsizes the inflammatory response of macrophages. Nature Materials 17, 1134–1144 (2018).

29. Okamoto, T. et al. Reduced substrate stiffness promotes M2-like macrophage activation and enhances peroxisome proliferator-activated receptor gamma expression. Experimental cell research 367, 264–273 (2018).

30. Garg, K., Pullen, N.A., Oskeritzian, C.A., Ryan, J.J. & Bowlin, G.L. Macrophage functional polarization (M1/M2) in response to varying fiber and pore dimensions of electrospun scaffolds. Biomaterials 34, 4439–4451 (2013).

31. Tong, S., Zhu, H. & Bao, G. Magnetic iron oxide nanoparticles for disease detection and therapy. Mater Today 31, 86–99 (2019).

32. Patel, N.R. et al. Cell elasticity determines macrophage function. PloS one 7, e41024 (2012).

33. Skinner, B.M. & Johnson, E.E.P. Nuclear morphologies: their diversity and functional relevance. Chromosoma 126, 195–212 (2017).

34. Blakney, A.K., Swartzlander, M.D. & Bryant, S.J. The effects of substrate stiffness on the in vitro activation of macrophages and in vivo host response to poly(ethylene glycol)-based hydrogels. Journal of biomedical materials research. Part A 100, 1375–1386 (2012).

35. Maruyama, K. et al. Cyclic Stretch Negatively Regulates IL-1β Secretion Through the Inhibition of NLRP3 Inflammasome Activation by Attenuating the AMP Kinase Pathway. Frontiers in physiology 9 (2018).

36. Ballotta, V., Driessen-Mol, A., Bouten, C.V.C. & Baaijens, F.P.T. Strain-dependent modulation of macrophage polarization within scaffolds. Biomaterials 35, 4919–4928 (2014).

37. Dziki, J.L. et al. The Effect of Mechanical Loading Upon Extracellular Matrix Bioscaffold-Mediated Skeletal Muscle Remodeling. Tissue engineering. Part A 24, 34–46 (2018).

38. McWhorter, F.Y., Wang, T., Nguyen, P., Chung, T. & Liu, W.F. Modulation of macrophage phenotype by cell shape. Proceedings of the National Academy of Sciences 110, 17253–17258 (2013).

39. Gao, Z., He, Q., Peng, B., Chiao, P.J. & Ye, J. Regulation of nuclear translocation of HDAC3 by IkappaBalpha is required for tumor necrosis factor inhibition of peroxisome proliferator-activated receptor gamma function. J Biol Chem 281, 4540–4547 (2006).

40. Jain, N., Iyer, K.V., Kumar, A. & Shivashankar, G.V. Cell geometric constraints induce modular gene-expression patterns via redistribution of HDAC3 regulated by actomyosin contractility. Proceedings of the National Academy of Sciences 110, 11349–11354 (2013).

41. Sun, S. et al. Monodisperse MFe2O4(M = Fe, Co, Mn) Nanoparticles. Journal of the American Chemical Society 126, 273–279 (2004).

42. Yang, H., Ogawa, T., Hasegawa, D. & Takahashi, M. Synthesis and magnetic properties of monodisperse magnetite nanocubes. Journal of Applied Physics 103, 07D526 (2008).

43. Tong, S., Hou, S., Ren, B., Zheng, Z. & Bao, G. Self-Assembly of Phospholipid–PEG Coating on Nanoparticles through Dual Solvent Exchange. Nano Letters 11, 3720–3726 (2011).

44. Tong, S., Quinto, C.A., Zhang, L., Mohindra, P. & Bao, G. Size-Dependent Heating of Magnetic Iron Oxide Nanoparticles. ACS nano 11, 6808–6816 (2017).

45. Mocarelli, P., Palmer, J. & Defendi, V. A permanent line of macrophages with normal activity in a primary antibody response in vitro. Immunol Commun 2, 441–447 (1973).

46. Qiu, Y. et al. Magnetic forces enable controlled drug delivery by disrupting endothelial cell-cell junctions. Nature Communications 8, 15594 (2017).

47. Qiu, Y. et al. Magnetic forces enable controlled drug delivery by disrupting endothelial cell-cell junctions. Nat Commun 8, 15594 (2017).

48. Mirza, R. & Koh, T.J. Dysregulation of monocyte/macrophage phenotype in wounds of diabetic mice. Cytokine 56, 256–264 (2011).

49. Furman, D. et al. Chronic inflammation in the etiology of disease across the life span. Nature Medicine 25, 1822–1832 (2019).

50. Yang, K. et al. Macrophage-mediated inflammatory response decreases mycobacterial survival in mouse MSCs by augmenting NO production. Scientific Reports 6, 27326 (2016).

51. Ferrer, J.M. et al. Measuring molecular rupture forces between single actin filaments and actin-binding proteins. Proceedings of the National Academy of Sciences 105, 9221–9226 (2008).

52. Kishino, A. & Yanagida, T. Force measurements by micromanipulation of a single actin filament by glass needles. Nature 334, 74–76 (1988).

53. Yu, L. et al. MRTF-A mediates LPS-induced pro-inflammatory transcription by interacting with the COMPASS complex. Journal of Cell Science 127, 4645–4657 (2014).

54. Leus, N.G. et al. HDAC 3-selective inhibitor RGFP966 demonstrates anti-inflammatory properties in RAW 264.7 macrophages and mouse precision-cut lung slices by attenuating NF-kappaB p65 transcriptional activity. Biochem Pharmacol 108, 58–74 (2016).

55. Scirpo, R. et al. Stimulation of nuclear receptor peroxisome proliferator–activated receptor‐γ limits NF‐κB‐dependent inflammation in mouse cystic fibrosis biliary epithelium. Hepatology 62, 1551–1562 (2015).

56. Lin, H. et al. Efficient in vivo bone formation by BMP-2 engineered human mesenchymal stem cells encapsulated in a projection stereolithographically fabricated hydrogel scaffold. Stem Cell Research & Therapy 10 (2019).

