## Supplement information for "Magnetic nanocomplexes coupled with an external magnetic field modulate macrophage phenotype – a non-invasive strategy for bone regeneration"

### 6. Supplementary information

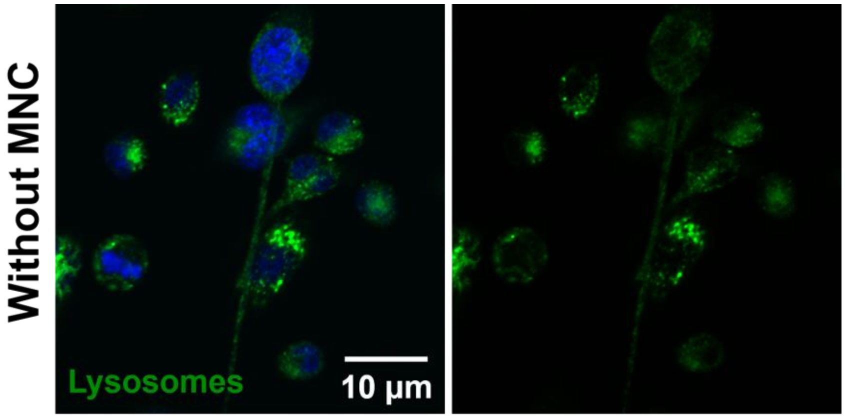


**Supplementary Fig.1:** Lysosome staining without MNC in macrophages. *Scale bar: 10 µm*

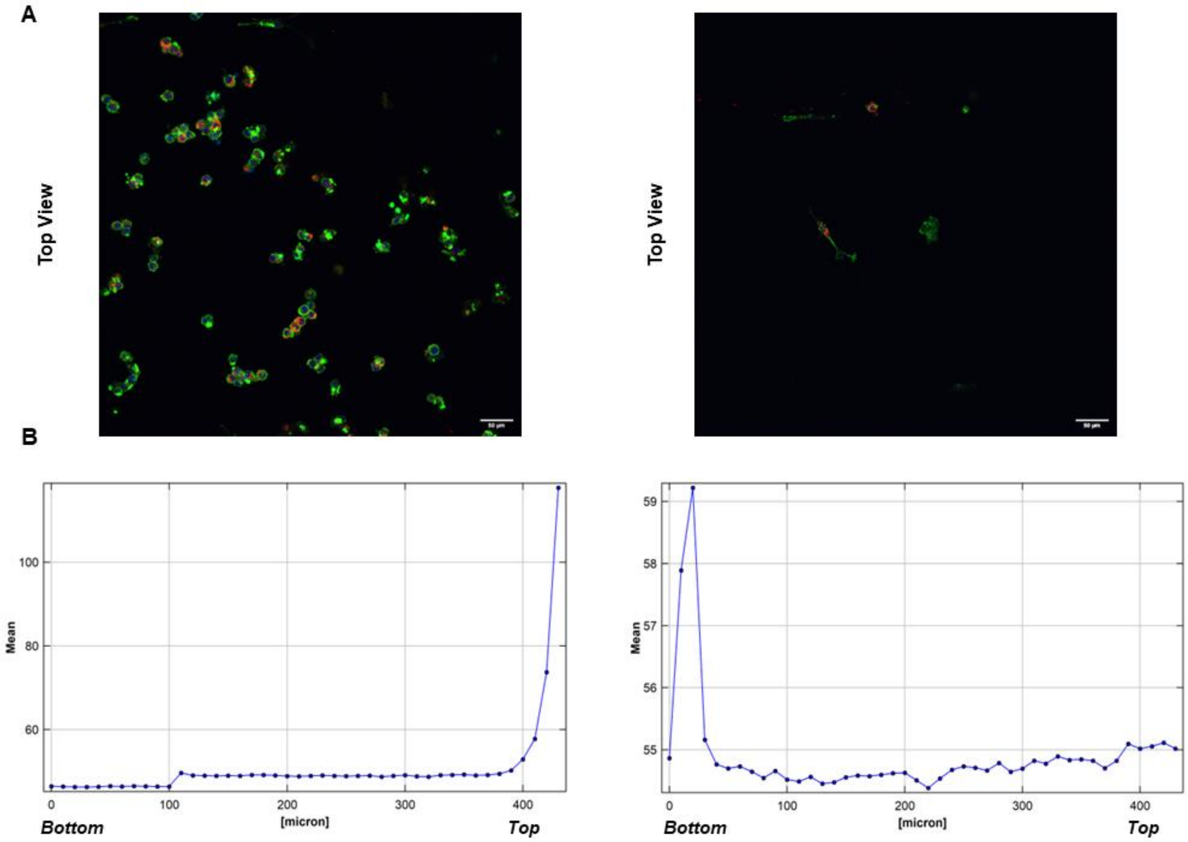


***Supplementary Fig.2: A)*** *M1-like macrophage loaded with MNC seeded on fibrin hydrogel without magnet (top left) and with the external magnet (top right) where 95% of the cells migrated through the fibrin hydrogel due to intracellular magnetic force. (F actin is stained in green and MNC in Red)* ***B)*** *Total fluorescent measured from top to bottom of the fibrin*

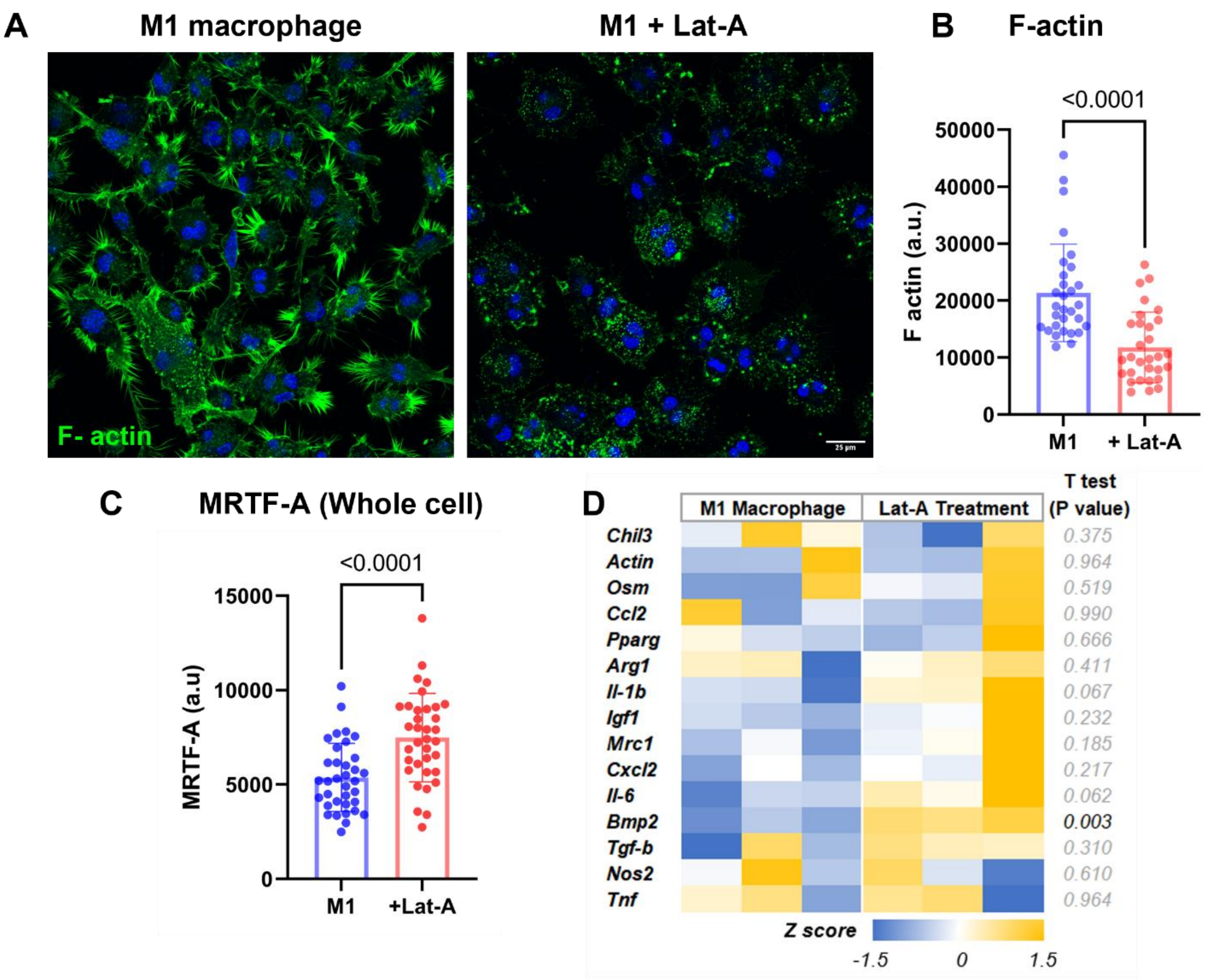


***Supplementary Fig.3: A)*** *M1-like macrophages treated with Lat-A stained for F actin in green.* ***B)*** *Quantification of F actin and* ***C)*** *whole cell MRTF-A in control and Lat-A treated macrophages.* ***D)*** *Heat map representing Z score of mRNA levels of inflammatory and anti-inflammatory genes in M1-like and with Lat-A treatment. Gapdh was used for normalization.*

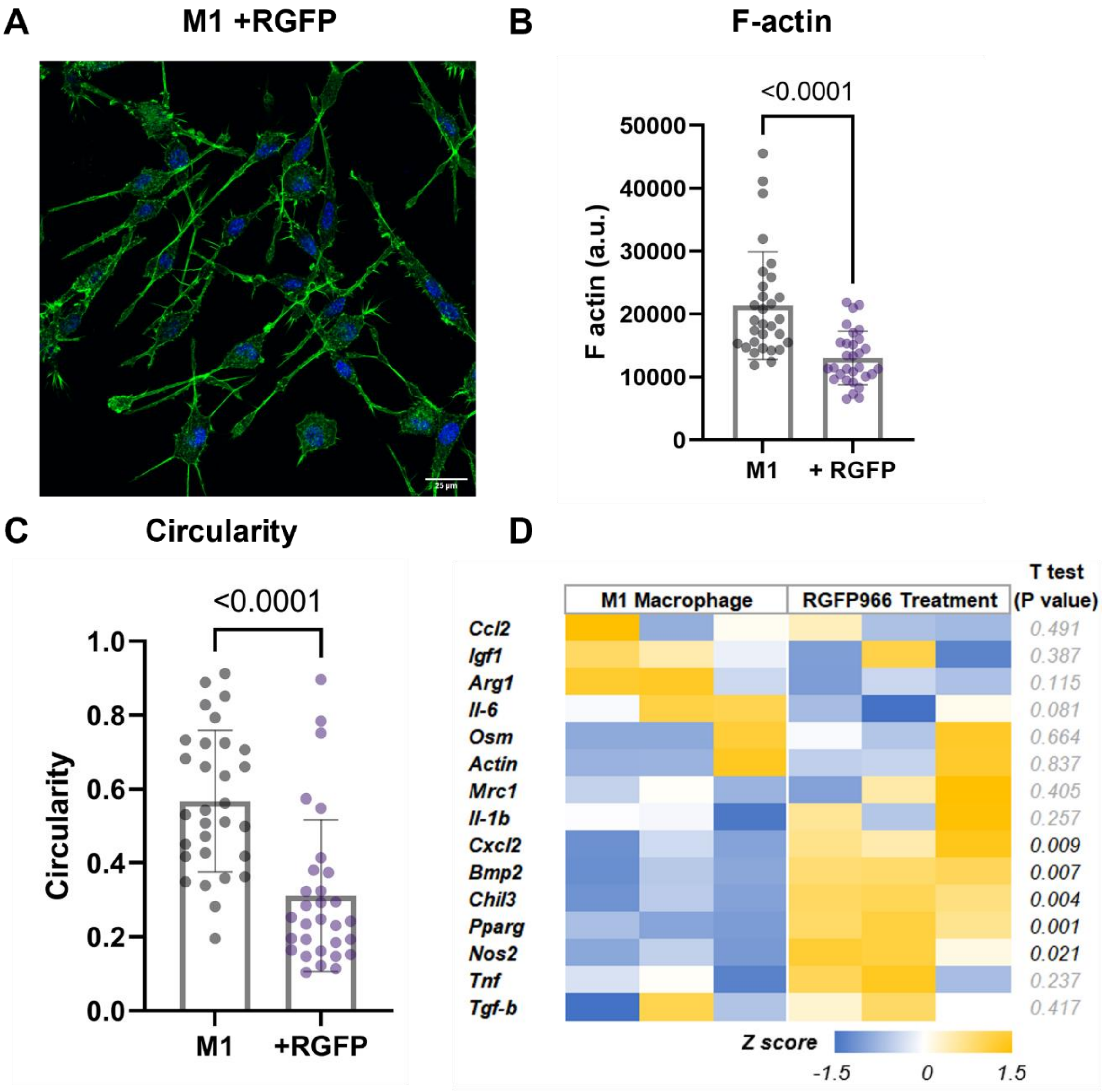


***Supplementary Fig.4: A)*** *M1-like macrophages treated with RGFP966 stained for F actin in green.* ***B)*** *Quantification of F actin and* ***C)*** *cell circularity in control and Lat-A treated macrophages.* ***D)*** *Heat map representing Z score of mRNA levels of inflammatory and anti-inflammatory genes in M1-like and with RGFP966 treatment. Gapdh was used for normalization.*

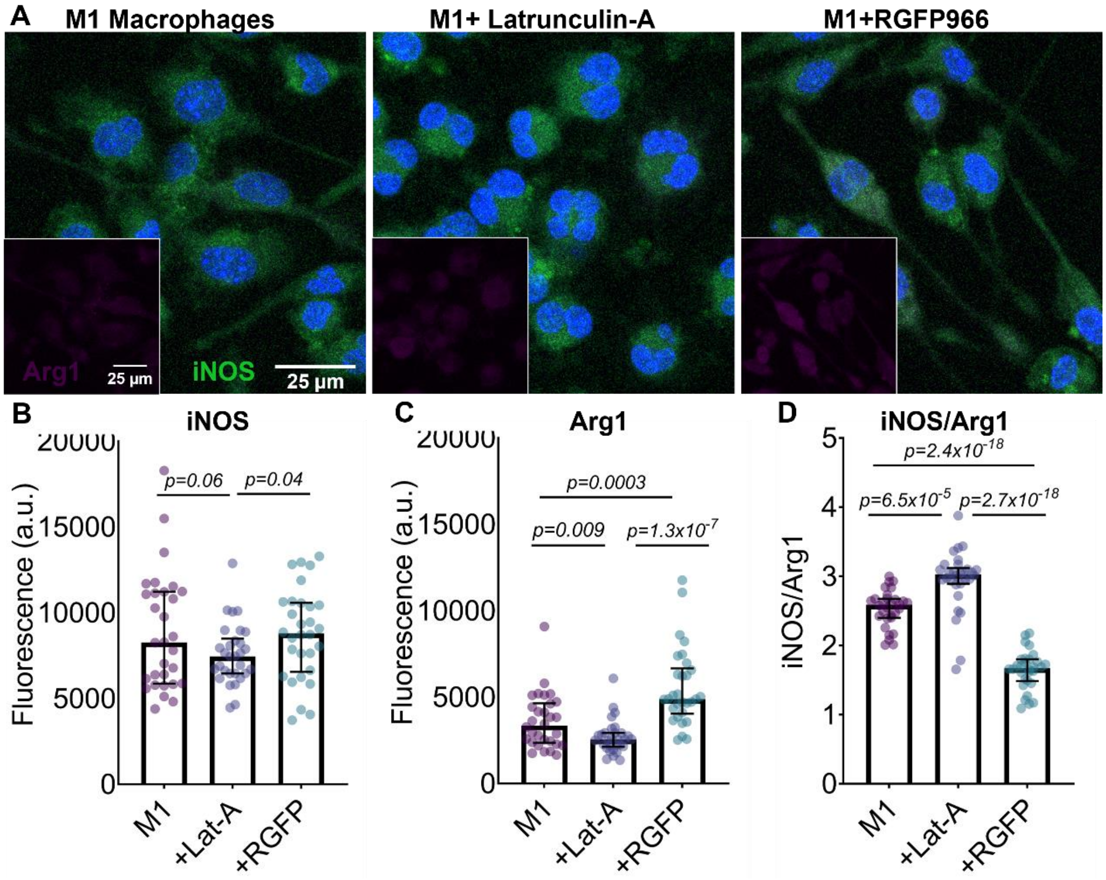


***Supplementary Fig.5: A)*** *M1-like macrophages treated with Lat-A and RGFP966 stained for iNOS in green and Arg1 in magenta.* ***B)*** *Quantification of iNOS and c) Arg1 in control, Lat-A and RGFP treated macrophages.*

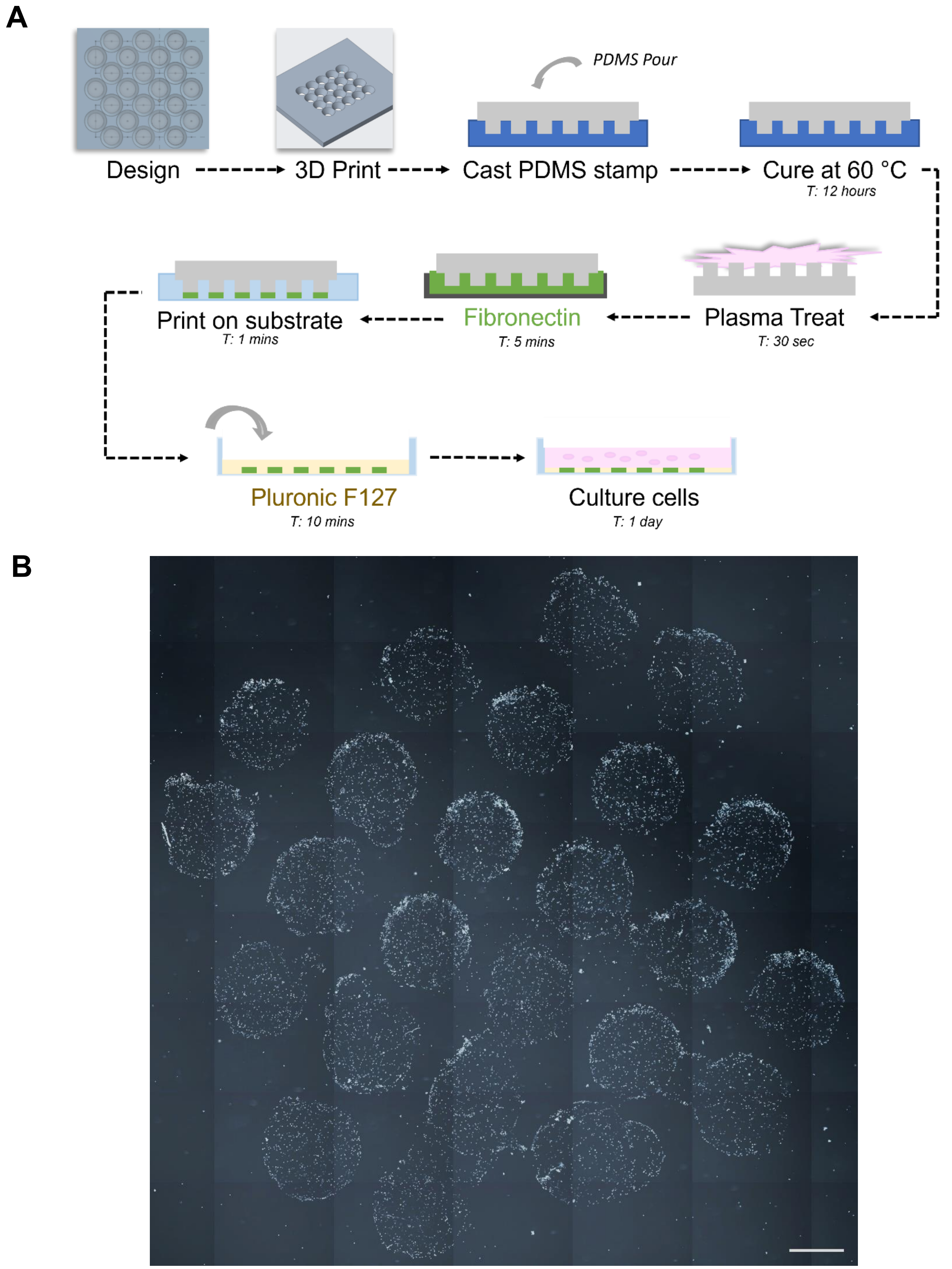


***Supplementary Fig.6 A)*** *Schematic showing sequential steps in the development of micropatterned substrate* ***B)*** *Macrophages attached to the fibronectin printed micropatterned substrate. Scale bar: 1 mm*
